# Protein-Free Catalysis of DNA Hydrolysis and Self-Integration by a Ribozyme

**DOI:** 10.1101/2024.08.31.610620

**Authors:** Deni Szokoli, Hannes Mutschler

## Abstract

Group II introns are ancient self-splicing ribozymes and retrotransposons. Though long speculated to have originated before translation, their dependence on intron-encoded proteins for splicing and mobility has cast doubt on this hypothesis. While some group II introns are known to retain part of their catalytic repertoire in the absence of protein cofactors, protein-free complete reverse splicing of a group II intron into a DNA target has never been demonstrated. Here, we demonstrate the complete independence of a group II intron from protein cofactors in all intron-catalyzed reactions. The ribozyme is capable of fully reverse splicing into single-stranded DNA targets *in vitro*, readily hydrolyzes DNA substrates, and is even able to unwind and react with stably duplexed DNA. Our findings make a protein-free origin for group II introns plausible by expanding their known catalytic capabilities beyond what would be needed to survive the transition from RNA to DNA genomes. Furthermore, the intron’s capacity to react with both single and double-stranded DNA in conjunction with its expanded sequence recognition may represent a promising starting point for the development of protein-free genomic editing tools.

## Introduction

Group II introns (G2Is) are among the largest and most complex naturally occurring catalytic RNAs (ribozymes) ever discovered, second only to the ribosomal RNAs (1, 2). They are categorized into three structural classes, IIA, IIB, and IIC, all of which catalyze the canonical self-splicing reaction based on two successive transesterification reactions, producing ligated exons and a 2’-5’ branched intron lariat (3). Most G2Is are mobile genetic elements and encode a signature protein with a dual reverse transcriptase (RT) and maturase function (4). Their mechanism of mobility, designated as “retrohoming”, is founded on the reversibility of the splicing reaction they catalyze (5). The reaction commences with the canonical self-splicing reaction of the G2I transcript, releasing an intron lariat (Figure 1A, steps 3, 2 and 1). The lariat is then capable of sequence-specifically binding to a DNA site within the host genome, where it inserts itself by reversing the self-splicing reaction with the bound DNA “exons” (Figure 1A, steps 1, 2 and 3), followed by reverse-transcription by the intron-encoded RT, which completes the G2I life cycle (6, 7).

**Figure 1:**
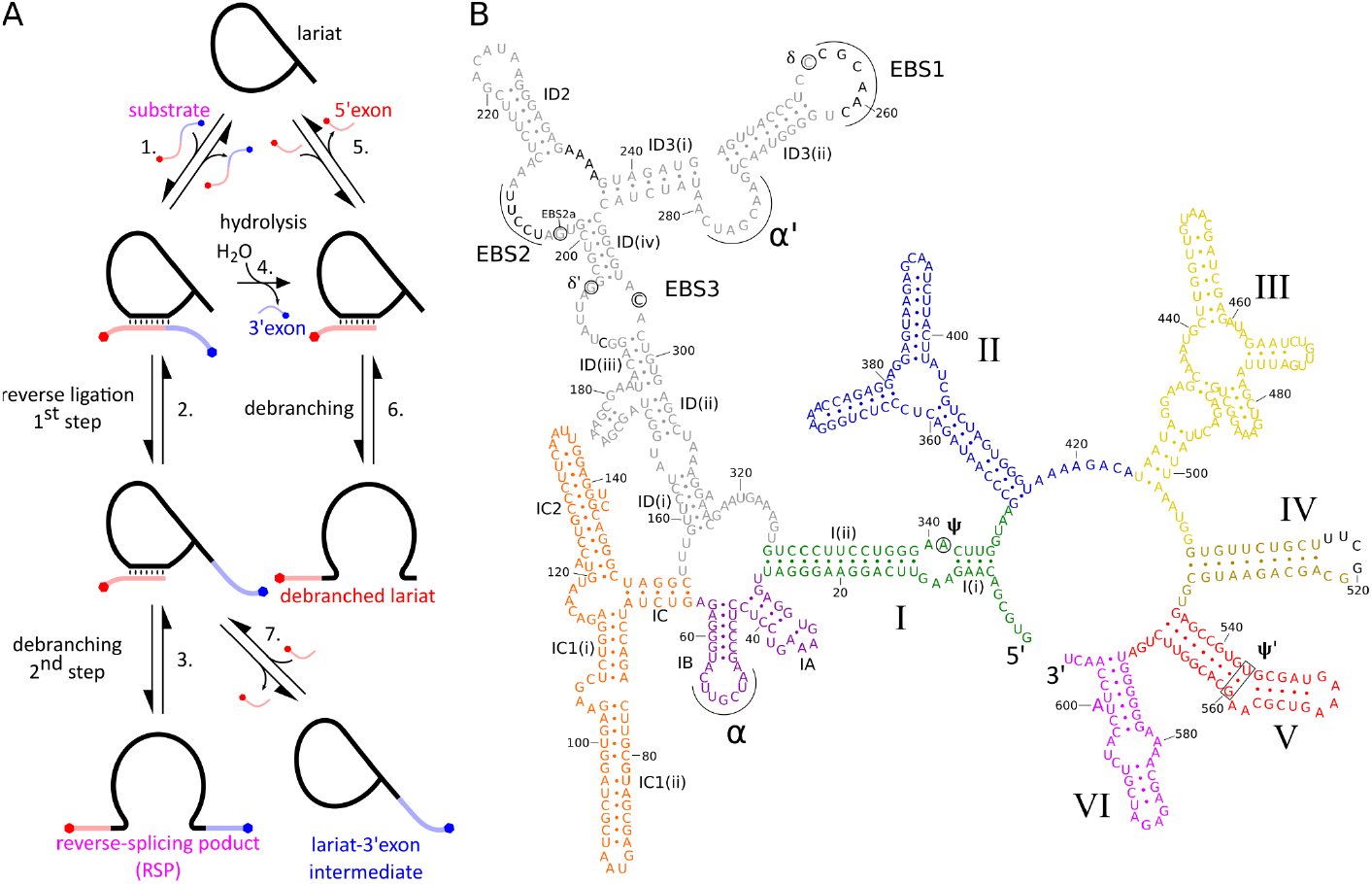
**A)** Schematic of the proposed reverse-splicing reaction network including all known sub-reactions. **B)** Secondary structure diagram of the modified group II intron P.li.LSU.I2. The sequence is based on the crystallization construct designed by Robart et al. (29). Letters in black font indicate mutations introduced in this study.

Although G2Is were initially investigated as model systems for pre-mRNA splicing, given their shared ancestry and catalytic mechanism with spliceosomes, the study of G2Is has subsequently yielded a range of highly effective biotechnological tools. The targetron system is the first retargetable RNA-guided genetic engineering system invented (8). Thermostable G2I encoded RTs have evolved to have a higher fidelity, processivity, and strand displacement activity than the previous industry standard retroviral RTs, likely due to a selection pressure to faithfully replicate their long and highly structured ribozymes (9, 10). A very recent addition to the G2I toolbox are the so-called “hydrolytic endonucleolytic ribozymes” (HYERs), which are ORF-less class IIC introns capable of sequence-specific ssDNA hydrolysis in mammalian cells (11). Though their class IIC ribozyme structure allows for a very limited sequence specificity, and they are unreactive with dsDNA, they show promise as a more compact and less immunogenic alternative to CRISPR-Cas-based gene editing systems (11, 12).

G2Is have co-evolved with their encoded proteins and are canonically thought of as ribonucleoprotein particles (13). One of the most conserved domains of the G2I IEP is the so-called “X”, or maturase domain, which binds tightly to the intron RNA, promotes its folding, and plays a part in the conformational changes involved in the splicing reaction (14– 16). The relationship between intron and IEP has become so important that many of the G2Is characterized are catalytically inactive in the absence of their IEP, only sometimes rescued by salt concentrations that far exceed physiological values (14, 17). The mechanism of G2I retrotransposition involves the reverse splicing of the intron lariat into a DNA site containing the intron binding sites (IBSs), followed by IEP-mediated reverse-transcription (6, 7). Previous research had shown that the model ORF-less intron bI1 found in the mitochondrial genome of *Saccharomyces cerevisiae*, was able to fully reverse splice into RNA targets, but only partially reverse splice into DNA targets (18, 19).

The discovery that introns with considerably reduced *in vitro* activity compared to bI1 are capable of complete reverse splicing into DNA with the assistance of the IEP (6, 15, 20), in conjunction with recent structural studies that have identified the IEP as a partial determinant of the correct positioning of the branchpoint adenosine, has led to the hypothesis that the IEP plays a crucial role in the reverse-splicing process (16, 21). In contrast, we suspected that the *status quo* assumption that G2I ribozymes require protein cofactors for the catalysis of any splicing reaction was shaped by sampling bias and early negative results (19, 22–26). We hypothesized that among the vast diversity of G2Is, some are capable of carrying out the full range of reactions on DNA substrates without the assistance of IEPs, allowing these ribozymes to be used as experimental test beds to study the ancient origin of G2Is in the context of pretranslational biology. Specifically, we found that a modified version of the class IIB1 self-splicing intron P.li.LSU.I2, encoded in the mitochondrial genome of the brown alga *Pylaiella littoralis* (27, 28), can catalyze full reverse self-splicing into DNA without any protein cofactor over a wide range of conditions. Our data refute the prevailing paradigm that IEPs are essential for the reverse splicing of G2Is into DNA and open up a new perspective on the evolutionary origin of G2Is, while raising the prospect of new biotechnological applications.

## Materials and methods

### DNA substrates and templates

All DNA oligos and gene blocks were procured from Integrated DNA Technologies. DNA substrates for reverse splicing were ordered with the 5’-Cy5 and 3’-6FAM labels and HPLC-purified by the manufacturer. Debranching DNA substrates were ordered with either 5’-Cy5 or 5’-6FAM labels and purified according to the manufacturer’s recommendation. When deemed necessary, the fluorescently labeled DNAs were PAGE purified by us.

The sequence for the modified P.li.LSU.I2 used in this study (Table S1C) was derived from the crystallization construct used by Robart *et al*. 2014 (29). The hairpin of domain IV was slightly shortened, and EBS 1, 2, and 3 were exchanged to be complementary to our DNA substrate. The secondary structure of the intron was then predicted by the UNAfold web server to ensure that the modifications did not disrupt the intron fold (30). The loops surrounding modified EBS2 and EBS3 were predicted to misfold, and compensatory mutations were introduced in order to restore the correct fold. The gene block was also designed to have a T7 RNA polymerase promoter at the 5’-end, followed by IBSs complementary to the new EBSs, as well as a short 3’-exon.

Templates for *in vitro* transcription were generated by PCR amplification of the gene block with a forward primer complementary to the T7 RNA polymerase promoter, and a reverse primer complementary to the 3’-exon. PCRs were performed in 100 µL reactions using Q5 Hot Start High-Fidelity 2X Master Mix (New England Biolabs - NEB), 0.5 µM of each primer, and 1 pmol of template. PCR products were purified using a Monarch PCR & DNA cleanup kit (NEB), and products were electrophoresed on an agarose TAE gel to confirm the product was of the correct size. Template concentration and purity were ascertained by absorbance at 260 nm using a Nanodrop spectrophotometer (Thermo Scientific).

### RNA preparation

Intron RNAs were transcribed *in vitro* with the TransciptAid T7 high yield transcription kit (Thermo Scientific), following the manufacturer’s instructions. The reactions were usually carried out overnight, and the modified P.li.LSU.I2 intron forward-spliced to a large degree within the IVT reaction buffer at 37°C. The DNA template was digested in the IVT buffer with the addition of Turbo DNAse (Thermo Scientific), according to the manufacturer’s instructions, and incubation for 30 minutes. One-third reaction volume of 4 M NH_4_Cl was added to a final concentration of 1 M, and the reaction temperature was increased to 47°C for another 30 minutes, to maximize forward-splicing. The samples were then purified with the Monarch RNA cleanup kit (NEB), and the RNA concentration was determined by absorbance at 260 nm on a Nanodrop spectrophotometer (Thermo Scientific). Lariat RNA was purified by electrophoresis on a 2 mm x 20 cm x 20 cm 6% 8 M urea PAGE (19:1 acrylamide:bisacrylamide), followed by gel extraction. Product bands were visualized by UV shadowing and excised with a sterile scalpel. Lariat RNA was recovered from the gel slices by the “crush and soak” method. Briefly, RNA was eluted from the crushed gel overnight at 4°C and constant agitation in ultrapure water, the supernatant was filtered off with a 0.22 µm cellulose acetate Spin-X filter (Corning) and precipitated in 0.3 M sodium acetate pH 5.2 and one volume of isopropanol. RNA pellets were washed with 80% ethanol, dried, and resuspended in nuclease-free water. Stock concentrations were determined by absorbance at 260 nm on a Nanodrop spectrophotometer (Thermo Scientific) and stored at −20°C.

### Reverse-splicing reactions

All reactions were prepared using a 10X stock solution of 400 mM Tris•HCl adjusted to pH 7.5 at room temperature. Standard reaction conditions for reverse splicing were 40 mM Tris•HCl pH 7.5, 0.001% PEG 8000, 0.5 M NH_4_Cl, 50 mM MgCl_2_, and varying concentrations of gel purified lariat and DNA substrates. Prior to initiating the reactions, the intron lariat was refolded in a buffer containing the Tris•HCl, PEG 8000, and the NH_4_Cl, by heating to 90°C for 1 minute, followed by slow cooling over the course of 10 minutes to the target reaction temperature, followed by another 10-minute incubation at the target temperature. The MgCl_2_ was then mixed in, followed by another 10-minute incubation at the target temperature, and then transferred to ice. Finally, the DNA substrate was added on ice and a sample was taken for the t = 0 time point before the remaining reaction samples were transferred to a thermocycler at the target temperature to initiate the reaction. Samples for gel electrophoresis were taken at increasing timepoints and quenched in nine volumes of denaturing gel-loading buffer (98% formamide, 10 mM EDTA pH 8, and a bromophenol blue tracking dye), taking care to always exceed the amount of MgCl_2_ with EDTA, and stored at −20°C. All experiments were performed in triplicate (n = 3). The DNA substrate used for all quantitative reverse-splicing experiments was “DNA RS sub. 1” (Table S1B). The DNA substrate used for reverse splicing which was later used as an input for RT-PCR and sequencing was “DNA RS sub. 2” (Table S1B).

The reverse-splicing reactions for screening optimal Mg^2+^ concentrations were performed according to the protocol noted above, except for some alterations in how the refolding and sampling were performed. For refolding, a master mix of intron lariat was heated, cooled, and then aliquoted into tubes containing varying MgCl_2_ concentrations. The entire 6 µl reactions were stopped after a one-hour incubation at 25°C, by quenching with 6 µl of 0.5 M EDTA pH 8 and 60 µl of denaturing gel loading buffer (98% formamide, 10 mM EDTA pH 8, and cresol red as a tracking dye), with an additional 0.2 µl of 0.5 M EDTA pH 8 in the samples containing 600 mM MgCl_2_.

The reverse-splicing reaction with the stable hairpin DNA substrate was performed according to the standard reaction protocol, except for differences in the final concentrations of some components (200 mM MgCl_2_, 1.5 µM lariat, and 0.1 µM hairpin DNA substrate). The reactions were carried out at 25°C overnight.

### Plasmid substrate preparation and reverse splicing

A minimized pGEM-T plasmid with the standard origin of replication and *ampR* gene was used as a backbone for cloning. Recombinant plasmids carrying a 50 bp insert containing either the intron IBSs or a stretch of dA_14_ in their place for the negative control, were generated by overhang PCR followed by ligation of the PCR blunt ends. Primers were designed to anneal to the plasmid with their 3’-ends while carrying 5’-overhangs corresponding to the two halves of the 50 bp insert. The resulting PCR amplicon was a linearized plasmid, flanked by the two halves of the 50 bp inserts. Blunt-ended PCR products were generated with the Q5 High-Fidelity 2X Master Mix (NEB), followed by 5’-phosphorylation, re-circularization, and DpnI digest with a KLD enzyme mix (NEB). The resulting 1894 bp recombinant plasmids were then transformed into chemically competent Top10 *E. coli* cells, which were grown overnight on LB plates containing carbenicillin (100 µg/mL). Clones carrying the expected recombinant plasmids were identified by Sanger sequencing performed by Microsynth. Recombinant plasmids were prepared from overnight cultures and purified with NucleoSpin Plasmid columns (Macherey-Nagel). The plasmids were linearized by restriction digestion with BspHI (NEB), for which the sole restriction site lies roughly equidistantly (949 bp downstream, and 941 bp upstream) from the intron insertion site in both directions. Linearized plasmids were purified using the Monarch DNA & PCR cleanup kit, and their concentration was determined by absorbance at 260 nm using a Nanodrop spectrophotometer (Thermo Scientific).

The procedure for the reverse-splicing reactions involving linearized plasmids followed the standard reverse-splicing protocol described in the previous section, except for some notable details. Reverse splicing into the plasmids was carried out at 25°C for 18.5 hours, under modified reaction conditions (40 mM Tris•HCl pH 7.5, 0.001% PEG 8000, 0.5 M NH_4_Cl, 150 mM MgCl_2_, 500 nM lariat, 7 nM linearized plasmid, and 1 U/µl Murine RNAse inhibitor (NEB)). The reaction was supplemented with RNAse inhibitor to reduce intron degradation from potential RNAse carryover from the plasmid miniprep. Meanwhile, the MgCl_2_ concentration and lariat:substrate ratio was increased with the hope of improving the detection of reverse-splicing products. Several controls were performed under identical reaction conditions, which included: 1) a control reaction with lariat and a plasmid lacking the IBSs; 2) only the lariat and no plasmid; 3) only the plasmid containing the IBSs and no lariat; 4) only the plasmid lacking the IBSs and no lariat.

### Reverse-transcription and RT-PCR

Because of the high salt concentrations in the reaction buffer, plasmid reverse-splicing samples were diluted 1:20 with nuclease-free water prior to reverse transcription. Reverse transcription was carried out using the Bst 3.0 DNA polymerase (NEB) because of its superior strand-displacement activity compared to other commercially available RTs (31), which we thought would be necessary to process the highly structured intron, as well as the dsDNA of the plasmid. Reverse transcriptions were carried out in 1X isothermal amplification buffer II, 6.25 mM MgSO_4_, 1 mM EDTA, 1.6 mM of each dNTP, 50 nM RT primer (“RS plasmid RT primer”, Table S1D), 1:200 dilution of the reverse-splicing sample and 0.64 U/µl Bst 3.0 DNA polymerase. A reaction tube containing nuclease-free water, the dNTPs, the RT primer, the reverse-splicing sample, and the EDTA was incubated at 65°C for 5 minutes, and then snap-cooled on ice for at least a minute, to facilitate the annealing of the RT primer to the intron. The EDTA was added at this step to chelate the Mg^2+^ carried over from the reverse-splicing reaction. In the following step, the isothermal amplification buffer II and MgSO_4_ were mixed in on ice, and a small aliquot was taken for the “no RT control” (NRC). Finally, Bst 3.0 was added to the RT reactions, and an equivalent amount of nuclease-free water was added to the NRC. Both the RT samples and the NRC were incubated at 65°C for 15 minutes to facilitate reverse transcription, after which the Bst 3.0 DNA polymerase was heat-inactivated at 80°C for 5 minutes.

RT-PCRs were performed on 1:100 diluted RT samples using hot start Taq DNA polymerase (NEB) supplemented with extreme thermostable ssDNA-binding protein (ET SSB, NEB), under the manufacturer’s recommended reaction conditions (1X standard Taq reaction buffer, 0.2 mM each dNTP, 0.2 µM each of “RS plasmid RT-PCR FWD” and “RS plasmid RT-PCR REV” (Table S1D), 25 U/ml hot start Taq DNA polymerase and 0.5 ng/µl ET SSB). Between the initial 1:200 dilution for the RT and the additional 1:100 dilution for the PCR, the initial splicing reactions containing 7 nM plasmid were diluted 20.000-fold down to 0.35 pM in the PCR reaction mix. Thermocycling was carried out with one cycle of 95°C for one minute, followed by 38 cycles of denaturing at 95°C for 15 seconds, annealing at 60°C for 15 seconds, and extension at 68°C for 15 seconds. The temperature ramp rate between the different steps was reduced to 4.5°C/second. RT-PCRs were performed in triplicate (n = 3). The RT-PCR products were electrophoresed on a 2% agarose TAE gel stained with SYBR safe (ThermoFisher Scientific). Prior to loading, 5 µl of each sample was mixed with 1 µl of 6X Purple Gel Loading Dye (NEB) and electrophoresed at 110 V for 30 minutes. The molecular weight standard used was the Quick-Load Purple 100 bp DNA Ladder (NEB). The PCR product band corresponding to the expected molecular weight of the RT-PCR product of the intron reverse spliced into the plasmid was excised from the gel, and the PCR product was extracted using the Monarch DNA gel extraction kit (NEB). The extracted PCR product was Sanger sequenced by Microsynth to confirm its identity.

Reverse transcription of the ssDNA (DNA RS sub. 2, Table S1B)) reverse-splicing product for Sanger sequencing was carried out in the same way as for the plasmid substrates, except that the template for the reaction was from a PAGE-purified product using the “ssDNA 3’SS RT primer” (Table S1D) to generate the 3’-splice-site cDNA, and “ssDNA 5’SS RT primer” to generate the 5’-splice-site cDNA. PCR of the RT product was done with the Q5 High-Fidelity 2X Master Mix (NEB), and two different primer pairs for the 5’- and 3’-splice-sites. The “ssDNA 3’SS RT-PCR REV” and “ssDNA 3’SS RT-PCR FWD” primers (Table S1D) were used for the 3’-splice-site, and the “ssDNA 5’SS RT-PCR FWD” and “ssDNA 5’SS RT-PCR REV” primers were used for the 5’ splice-site.

### Denaturing polyacrylamide gel electrophoresis

All gel mixes were prepared by mixing the desired ratio of stock solutions containing 8 M urea, 1X TBE, and either 20% or 0% acrylamide. Gel casting stock solutions contained a 19:1 ratio of acrylamide:bisacrylamide and were filtered and degassed. Gel extractions were performed using 6% gels (20 cm x 20 cm x 0.2 cm). PAGE of all reverse-splicing experiments was conducted using two-layer 5%/12% gels (20 cm x 20 cm x 0.1 cm). Two-layer gels were cast in two steps. First, the bottom half is cast by pouring a 12% acrylamide gel mix, and carefully layering 2 ml of isopropanol on top of the gel mix before it sets. Once the bottom layer is set, the isopropanol is decanted from the gel plates. Finally, the top half was cast using a 5% gel mix.

Samples in formamide loading buffer were denatured by heating to 95°C for 3 minutes, and directly transferred to ice until they were loaded onto the gel. The conductive media used for PAGE was 1X TBE. Electrophoresis of 1 mm thick gels would proceed at a constant 35 W until the bromophenol blue would reach the last two centimeters of the gel. All polyacrylamide gels were imaged using the Sapphire biomolecular imager (Azure Biosystems).

### Data analysis

Scans taken of PAGE gels were analyzed using the Azurespot image analysis software (Azure Biosystems). Only Cy5 fluorescence values were used for quantification due to a pH sensitivity issue with the 6FAM fluorophore. During electrophoresis, a pH gradient formed in the two-layer gels, leading to inaccurate 6FAM readings, particularly overreporting high-MW and underreporting low-MW band fluorescence, while Cy5 remained unaffected.

Bands corresponding to different molecular species were selected, and the backgrounds were subtracted. The band percentages (%Band) were exported and converted into molar concentration (C_species_) using the following equation:

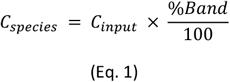

Where C_input_ is the initial concentration of substrate. Time course data was plotted and analyzed using GraphPad Prism 10 (Graphpad Software, Inc.). DNA substrate depletion displayed monophasic kinetic behavior and could be fit with a single exponential decay:

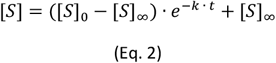

Where [*S*] is the molar concentration of the unreacted substrate at any timepoint *t*, [*S*]_0_ is the initial molar concentration of substrate at *t* = 0, and [*S*]_∞_ is the molar concentration of substrate as *t* approaches infinity.

Both the debranching and the accumulation of the free 5’-exon as a result of reverse splicing and DNA hydrolysis also displayed a monophasic kinetic behavior. The data were fitted to a single exponential:

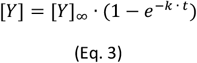

Where [*Y*] is the molar concentration of either the free 5’-exon or the debranched intron at any timepoint *t*, and [*Y*]_∞_ is the value of [*Y*] as *t* approaches infinity.

The reverse-splicing reaction displayed a biphasic kinetic behavior and was fitted with a double-exponential:

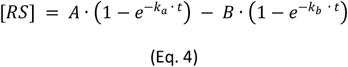

Where [*RS*] is the molar concentration of the reverse-splicing product at any timepoint *t, A* and *B* and *k*_*a*_ and *k*_*b*_ are the amplitudes and apparent rate constants of the first and second reaction phase, respectively.

The Cy5 fluorophore used for quantification underwent an unexpected bleaching in the 600 mM MgCl_2_ condition of the Mg^2+^ titration experiment (Figure 2A). Therefore, the percentage of reverse-splicing product (RSP) for the 600 mM MgCl_2_ condition was calculated by correcting the values obtained from the corresponding 6FAM fluorescence of the same band. A gel-specific correction factor was calculated by dividing the volume of each RSP band in the Cy5 channel by the corresponding volume of the same band in the FAM channel for each gel, and averaging all the ratios obtained this way. The corrected Cy5 RSP percentages for the bands in the 600 mM MgCl_2_ condition were calculated by multiplying the volume of the band obtained from the FAM channel with the gel-specific correction factor.

**Figure 2:**
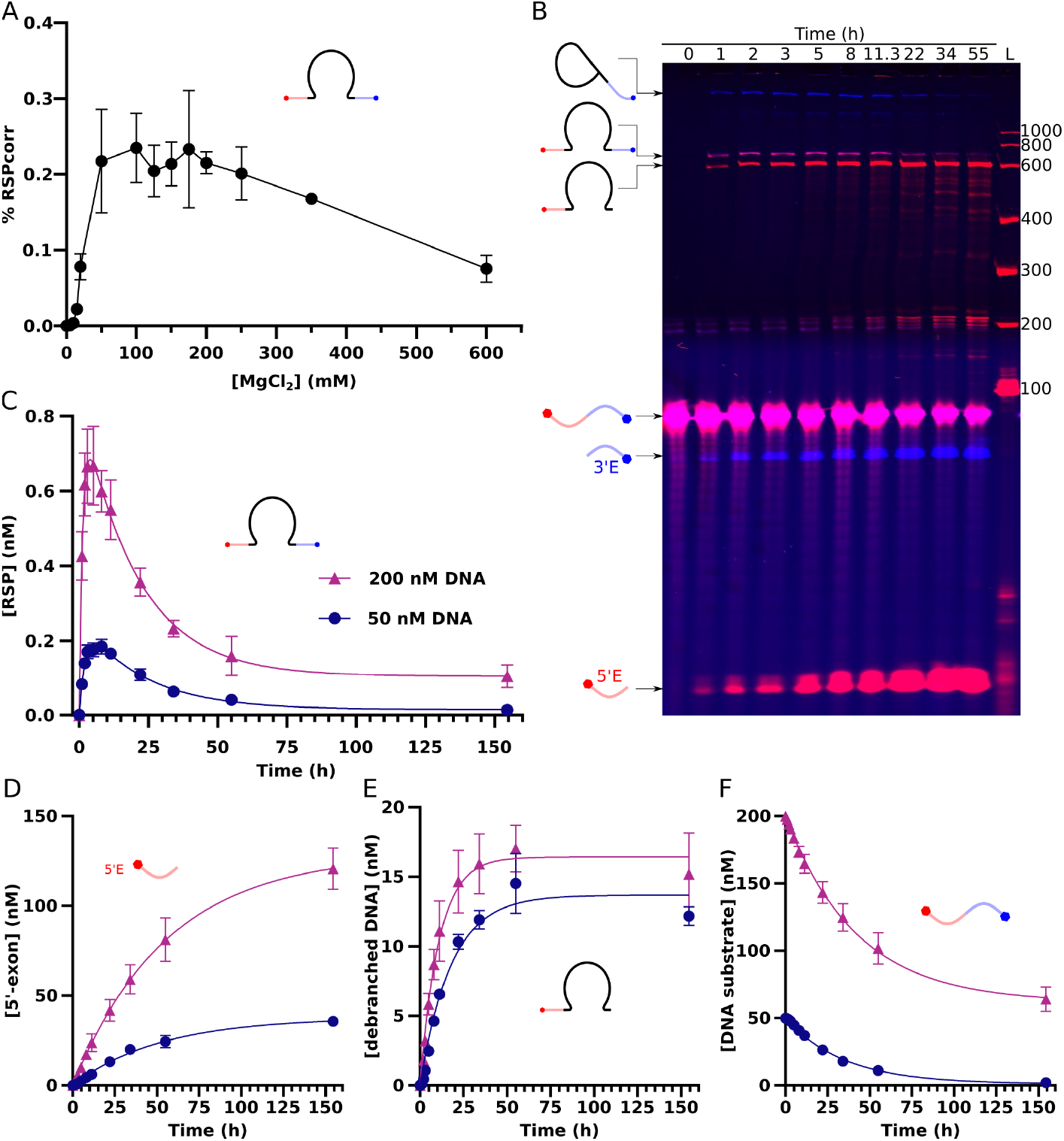
Biochemical characterization of protein-free reverse-splicing into single-stranded DNA. **A)** Percentage of substrate converted to RSP after one hour of incubation at 25°C of 100 nM intron lariat and 200 nM DNA substrate, in a buffer containing 40 mM Tris•HCl pH 7.5, 0.5 M NH_4_Cl, 0.001% PEG 8000, and varying concentrations of MgCl_2_. Values were corrected for the 600 mM MgCl_2_ sample (see data analysis section in materials and methods). **B)** Example denaturing PAGE of the reverse-splicing reaction time course. Conditions are the same as in A), except the reaction was carried out at 30°C and in 50 mM MgCl_2_. Bands are labeled with a drawing of the corresponding species. **C), D), E)**, and **F)** are kinetic traces of RSP, free 5’-exon, debranching product, and the DNA substrate respectively. Conditions as in B), except that those reactions represented by purple triangles contained 200 nM DNA substrate, and those with dark blue circles contained 50 nM DNA substrate. Datapoints are the averages of triplicates. Error bars represent ± standard deviations (n = 3).

The first dataset (n = 3) collected for the Mg^2+^ titration plot (Figure 2A) had a high error in the range between 100 mM and 200 mM MgCl_2_. In an attempt to better characterize the reaction in this range the experiments were repeated for the 50, 100, 125, 150, 175, 200, 250, and 600 mM MgCl_2_ conditions (n = 3). The combined means for each condition were calculated with the following equation:

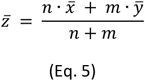

Where 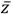 is the combined sample mean, *n* and *m* are the numbers of samples in each set, and 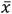 and 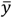 are the means of each set of samples. The sample standard deviation for each experiment was calculated using the following equation:

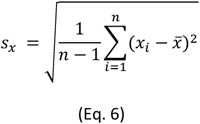

The combined standard deviation of both datasets 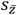 was calculated using the following equation:

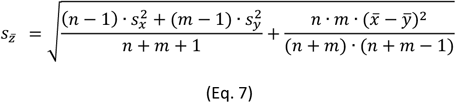

The points plotted on Figure 2A represent the combined means 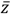 of the two datasets for each condition, and the error bars represent the combined standard deviation 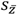.

## Results

### The group II intron P.li.LSU.I2 is capable of protein-free reverse splicing into single-stranded DNA

To ascertain the capacity of group II introns to fully reverse splice into DNA without the aid of proteins, we initially handpicked two group II introns from the literature for an initial screen of protein-free DNA reverse splicing. We avoided ribozymes known to be unreactive with DNA substrates and focused primarily on the ones that either had a published 3D structure or were known to autocatalytically self-splice *in vitro* at low Mg^2+^ concentrations (19, 22, 29, 32), which was taken as a sign that protein cofactors might not be necessary for proper folding and catalysis. The chosen introns according to these criteria were the model intron P.li.LSU.I2, which interrupts the mitochondrial large subunit rRNA gene in the brown alga *Pylaiella littoralis* (27, 28), and C.i.SSU.I1, a recently discovered highly reactive intron that interrupts the small subunit rRNA gene in the pathogenic fungi *Coccidioides immitis* (32).

Next, we designed a suitable DNA substrate to assay partial and full reverse splicing (Figure 1A). The substrate DNA was designed to contain intron binding sites (IBS) and a Cy5 label at its 5’ terminus, as well as a FAM label at its 3’ terminus. The two fluorophores allowed for the orthogonal detection of reaction products of both full and partial reverse splicing (Figure 1A). The long ORF contained in domain IV (D4) of both introns was replaced with short loops. The sequence of C.i.SSU.I1 was left otherwise unchanged, as its wild-type IBSs were used in the design of the DNA substrate. In contrast, several mutations were introduced into the exon binding sites (EBS) of P.li.LSU.I2 so that they would be complementary to the substrate DNA, in addition to several compensatory mutations in nearby regions which maintained the predicted fold of the intron (Table S1A, Figure 1B).

We then performed a pilot reverse-splicing assay of the selected intron lariats. We found both introns to be capable of reversing the second step of forward-splicing on a DNA substrate (Figure S1), which agrees with previous reports (19). To our surprise, we found that the modified P.li.LSU.I2 appeared to also be capable of the more elusive reaction of debranching with the DNA substrate (Figure S1) since we observed the emergence of a P.li.LSU.I2 reaction product carrying both fluorophores, a reaction previously demonstrated only with chimeric RNA/DNA substrates, or with the aid of proteins (6, 22, 33). In contrast, an equivalent doubly labelled product could not be observed in reactions carried out with C.i.SSU.I1. To determine the identity of the P.li.LSU.I2 reaction product carrying both fluorophores we reverse-transcribed and amplified the gel-extracted band using primers specific for the 5’- and 3’-splice-sites. Subsequent Sanger sequencing revealed the intron to be sequence-specifically inserted at the expected site between EBS1 and EBS3, thus confirming complete reverse splicing into ssDNA (Figure S2). Based on this, the P.li.LSU.I2 intron was chosen for further characterization of the protein-free reverse-splicing reaction.

### Biochemical characterization of intron reverse splicing into DNA

Studying complete reverse splicing is challenging due to the potential co-occurrence of undesired side-reactions. The reverse-splicing reaction is carried out in two sequential, fully reversible transesterification steps. The first step is reverse ligation (Figure 1A, step 2), where the 3’-OH of the lariat performs a nucleophilic attack on the phosphate belonging to the IBS3 nucleotide of the ligated exons, producing the lariat-3’-exon intermediate, and a free 5’-exon. This is followed by the second step – debranching (Figure 1A, step 3), where the 3’-OH of the 5’-exon performs a nucleophilic attack on the phosphate in the branch site of the lariat-3’-exon intermediate, producing a linear intron with covalently linked 5’- and 3’-exons on either end – the reverse-splicing product (RSP). We expected that some intron lariats would be capable of debranching with the 5’-exons (Figure 1A, steps 5 and 6) released by the incomplete reverse splicing of other lariat-3’-exon intermediates (Figure 1A, step 7).

Considering that these reactions are reversible, it would be reasonable to assume that the RSP and debranched lariat concentrations would monotonically approach an equilibrium concentration given sufficient time. However, based on the ubiquitous accumulation of free 3’-exon throughout reverse-splicing experiments (e.g. Figure 2B), which is not a product of the reverse-splicing reactions considered thus far, we postulated the occurrence of an additional reaction, analogous to the spliced exon reopening (SER) reaction. This reaction canonically involves a group II intron hydrolytically cleaving an RNA substrate containing the sequence of its ligated exons (34). The key difference between canonical SER and the reaction observed here is that the substrate is not an RNA, but a single-stranded DNA (Figure 1A, step 4).

To explore the full reverse-splicing activity of P.li.LSU.I2 in more depth, the DNA substrate was reacted with the purified lariat at 25°C for one hour in 0.5 M NH_4_Cl and varying concentrations of MgCl_2_. Interestingly, under these conditions, the yield of fully reverse-spliced DNA peaked after 50 mM Mg^2+^, followed by a decrease, instead of a plateau at Mg^2+^ concentrations above 250 mM Mg^2+^ (Figure 2A). This was in contrast to the other reaction products such as the debranched intron and the free 5’-exon, whose fraction increased even at higher Mg^2+^ concentrations (Figure S3). This indicated that the observed decrease in RSP was not due to an overall inhibition of catalysis at higher Mg^2+^ concentrations, but rather the result of the DNA hydrolysis side-reaction that increasingly dominates the reaction outcome at high Mg^2+^ concentrations. In agreement with this explanation, time-course experiments of reverse splicing conducted at 50 mM and 350 mM MgCl_2_ suggested that a greater increase in the rate of RSP depletion compared to that of RSP generation was responsible for the observed effect (Figure S4).

We performed a more detailed kinetic analysis of the reverse-splicing reaction at conditions that permitted convenient reaction times and only showed moderate RNA degradation (100 nM lariat, 40 mM Tris•HCl pH 7.5, 0.5 M NH_4_Cl, 50 mM MgCl_2_, 0.001% PEG 8000, at 30°C) at a 2:1 (200 nM DNA) and 1:2 (50 nM DNA) lariat to ssDNA substrate ratios. Under both conditions, the formation of RSP was characterized by an initial burst phase during the first ∼3 h of the reaction (t_1/2_ = 1.5 h for 50 nM DNA, and t_1/2_ = 0.87 h for 200 nM DNA, Table 1B). Over long timescales the quasi-irreversible hydrolysis of the DNA substrate (Figure 2B and 2D) results in a gradual decline in RSP levels (Figure 2C, t_1/2_ = 15.6 h for 50 nM DNA, and t_1/2_ = 14.1 h for 200 nM DNA), similar to previous reports of reverse splicing into RNA targets (35). It can be inferred from these results that P.li.LSU.I2 seems to catalyze efficient DNA hydrolysis (Figure 2B and 2D). This conclusion is supported by a recent publication that has independently found that certain class IIC introns are also capable of DNA hydrolysis (11).

**Table 1:** Kinetic parameters from biochemical characterizations.

**A) Complete reverse splicing – monophasic kinetics:**
| species | DNA substrate (nM) | [Y]∞ or [S]∞ (nM) [95% CI] | k (h<sup>-1</sup>) [95% CI] | t<sub>1/2</sub> (h) [95% CI] |
| --- | --- | --- | --- | --- |
| debranched product – [Y] | 50 | 13.7 [12.8, 14.6] | 0.056 [0.047, 0.066] | 12 [10, 14] |
| debranched product – [Y] | 200 | 16 [15, 18] | 0.091 [0.076, 0.111] | 8 [6, 9] |
| free 5'-exon – [Y] | 50 | 39 [36, 40] | 0.019 [0.017, 0.022] | 36 [32, 41] |
| free 5'-exon – [Y] | 200 | 129 [120, 139] | 0.018 [0.015, 0.020] | 39 [34, 45] |
| substrate – [S] | 50 | 0* | 0.028 [0.025, 0.030] | 25 [24, 26] |
| substrate – [S] | 200 | 62 [53, 69] | 0.024 [0.021, 0.027] | 29 [26, 33] |

\* The [S] ∞ parameter was manually set to 0.

**B) Complete reverse splicing – biphasic kinetics:**
| species | DNA substrate (nM) | A [95% CI] | k<sub>A</sub> (h<sup>-1</sup>) [95% CI] | t<sub>1/2 A</sub> (h) [95% CI] | B [95% CI] | k<sub>B</sub> (h<sup>-1</sup>) [95% CI] | t<sub>1/2 B</sub> (h) [95% CI] |
| --- | --- | --- | --- | --- | --- | --- | --- |
| RSP – [RS] | 50 | 0.26 [0.23, 0.31] | 0.46 [0.37, 0.57] | 1.5 [1.2, 1.9] | -0.28 [-0.29, -0.22] | 0.04 [0.03, 0.06] | 16 [12, 20] |
| RSP – [RS] | 200 | 0.85 [0.76, 1.00] | 0.8 [0.61, 1.05] | 0.87 [0.66, 1.15] | -0.75 [-0.87, -0.65] | 0.05 [0.03, 0.07] | 14 [10, 20] |

**C) Isolated biphasic debranching kinetics:**
| [D]∞ (nM) [95% CI] | F<sub>fast</sub> [95% CI] | k<sub>fast</sub> (h<sup>-1</sup>) [95% CI] | t<sub>1/2 fast</sub> (h) [95% CI] | F<sub>slow</sub> [95% CI] | k<sub>slow</sub> (h<sup>-1</sup>) [95% CI] | t<sub>1/2 slow</sub> (h) [95% CI] |
| --- | --- | --- | --- | --- | --- | --- |
| 75 [70, 98] | 0.3 [0.2, 0.5] | 10 [5, 19] | 0.07 [0.04, 0.13] | 0.7 [0.5, 0.8] | 1.3 [0.4, 2.1] | 0.5 [0.3, 1.8] |

In contrast to protein-free reverse-ligation (19) (Figure 1A, step 2), protein-free debranching of DNA substrates (Figure 1A, steps 3 and 6) by group II intron ribozymes has not been previously documented, to the best of our knowledge. By incubating the intron lariat with free 5’-exon DNA we were able to study debranching (Figure 1A, step 6) in the absence of reverse-ligation (Figure 1A, step 2) and hydrolysis (Figure 1A, step 4). Under the employed reaction conditions (40 mM Tris•HCl pH 7.5, 0.1% Tween 20, 0.5 M NH_4_Cl, 50 mM MgCl_2_, 100 nM lariat, and 1 µM 5’-exon DNA at 25°C) formation of the debranched product was characterized by a biphasic kinetic behavior, with an apparent half-life of 4.3 min for the fast phase, and 31 min for the slow phase (Table 1C, Figure S5). The biphasic kinetics are likely caused by two populations of lariat: one that is bound to the substrate at t = 0, and one that is unbound. The substrate-bound population reacts with fast kinetics, while the unbound population must first bind to the substrate before reacting, resulting in an apparent slower reaction phase.

### P.li.LSU.I2 is capable of processing double-stranded DNA

Having found that reverse splicing into single-stranded DNA can occur without the involvement of proteins emboldened us to consider whether there might be additional functions typically attributed to proteins for which the ribozyme alone might be sufficient. In particular, we wondered if a protein-free P.li.LSU.I2 ribozyme alone was capable of processing dsDNA substrates, i.e. catalyzing its own insertion into a hybridized target strand. In the subclasses of group II introns that retrohome into dsDNA (like in the case of P.li.LSU.I2), it is generally understood that the IEP is responsible for dsDNA binding and unwinding (36, 37).

With the goal of ensuring a 1:1 stoichiometry of DNA strands during our experiments, we cloned the modified P.li.LSU.I2 IBSs into a plasmid derived from the standard vector pGEM-T. The plasmid was subsequently linearized by restriction digest with the endonuclease BspHI, with a single restriction site positioned roughly equidistantly from either end of the IBSs (see materials and methods). The result was a 1.9 kb dsDNA with the intron insertion site flanked on either end with ∼950 bp (Table S1B). Because of the anticipated low efficiency of the reaction and lack of fluorescent labeling, we utilized an RT-PCR-based method to detect full reverse splicing (Figure 3A). After incubating 7 nM of the dsDNA substrate with 0.5 µM of purified P.li.LSU.I2 lariat for 18.5 h at 25 °C, the reaction products were reverse-transcribed with a primer specific to the intron, followed by PCR with a reverse primer specific to the intron, and a forward primer specific to the plasmid (Figure 3B). Strikingly, we were able to detect an RT-PCR product of the expected molecular weight, suggestive of covalent integration of P.li.LSU.I2 into the dsDNA target (Figure 3C). This notion was further supported by the absence of any PCR product in the absence of either the P.li.LSU.I2 lariat or the reverse transcriptase. Furthermore, no RT-PCR product was observed after incubation of the lariat with a control plasmid devoid of the P.li.LSU.I2 IBSs (Figure 3C).

**Figure 3:**
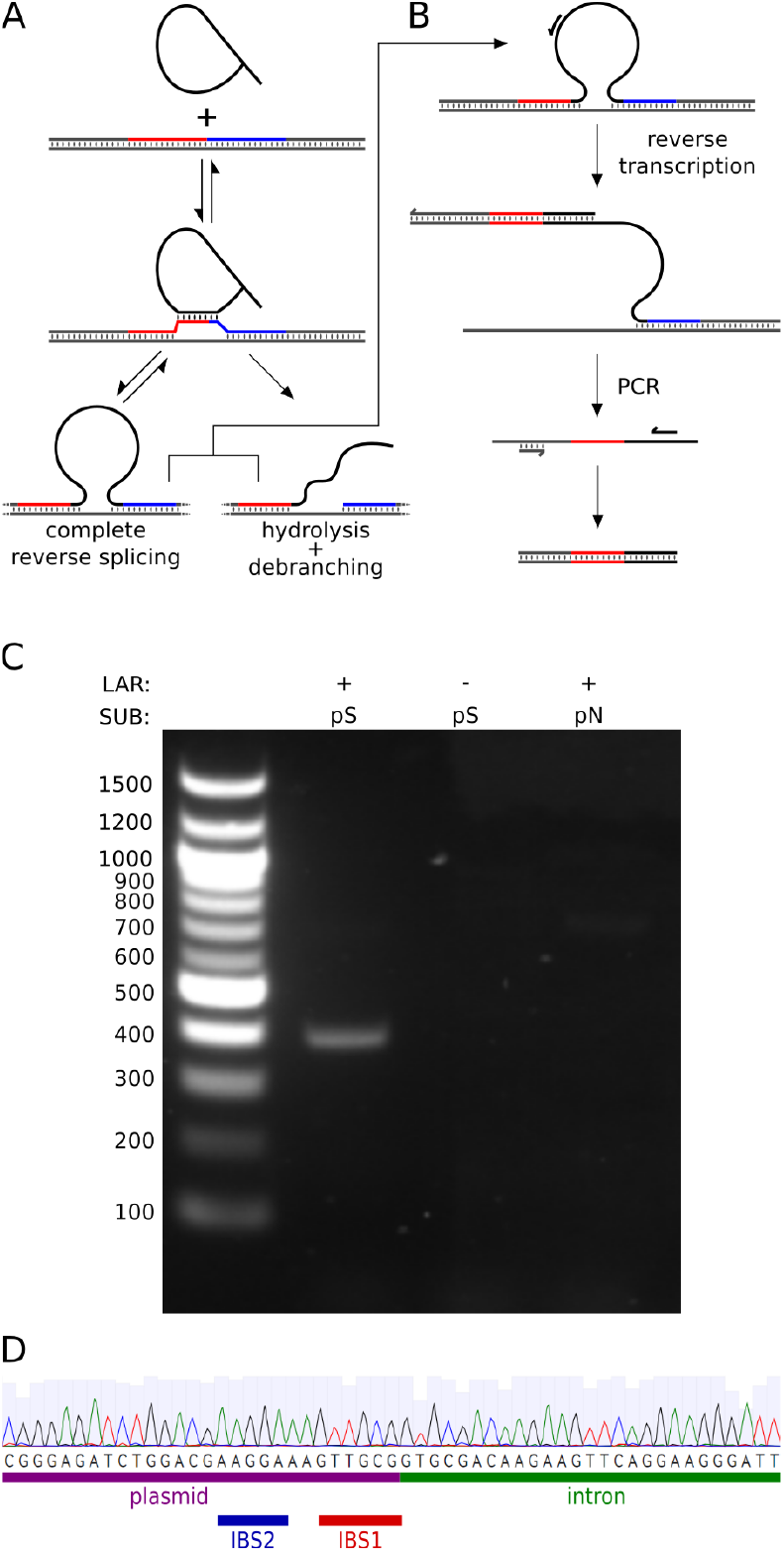
**A)** Reaction scheme of P.li.LSU.I2 reverse splicing into a dsDNA. The intron unwinds the dsDNA, base pairs with the IBSs, and catalyzes reverse splicing and hydrolysis of DNA. **B)** For analysis, the reaction products were reverse-transcribed with a primer specific to the intron, followed by a PCR with a reverse primer specific to the intron, and a forward primer specific to the plasmid DNA. **C)** 2% agarose TAE gel of RT-PCRs in absence or presence of lariat (LAR), and plasmid substrate (SUB) either with (pS) or without (pN) the intron insertion site. **D)** Section of the Sanger sequencing chromatogram of the RT-PCR product from C) showing that the intron is sequence-specifically inserted at the expected 5’-splice-site.

Ultimately, we could confirm integration of the ribozyme by Sanger sequencing of the RT-PCR product, which revealed that the product contained both the expected plasmid and intron sequences, and that the intron was inserted sequence-specifically at the intended site (Figure 3D). Due to the long incubation time required for dsDNA processing, we limited ourselves to detecting the ligated hybrid between the 5’-exon DNA target and P.li.LSU.I2, as we expected the DNA hydrolysis and 5’-exon debranching pathway (Figure 1A, steps 4 and 6) to dominate the product pool, while any potential full reverse-splicing product had already been lost after 18.5 h of incubation.

Having demonstrated that the intron can react with dsDNA we aimed to confirm that the hydrolysis / debranching reaction was indeed the dominant reaction trajectory using a gel-based assay. To this end, we designed a highly stable 89 nt fluorescently labeled DNA hairpin that contained the IBSs in the middle of the 43 bp double-stranded region (Figure 4A), which allowed for direct measurements of the reaction using PAGE. After a 20-hour incubation at an elevated Mg^2+^ concentration (1.5 µM lariat, 100 nM fluorescent DNA hairpin substrate, 40 mM Tris•HCl pH 7.5, 0.5 M NH_4_Cl, 200 mM MgCl_2_, 0.001% PEG 8000, at 25°C), we were able to detect a band at the expected molecular weight of the reverse-splicing product (Figure 4B). Notably, the product band seemed to only carry the Cy5 fluorescent label corresponding to the 5’-exon, while fluorescence from the 3’-exon FAM label was not detected. This finding was well in agreement with the RT-PCR-based assay used for the plasmid-based construct and confirmed that the lariat was capable of hydrolyzing, and subsequently debranching with dsDNA substrates.

**Figure 4:**
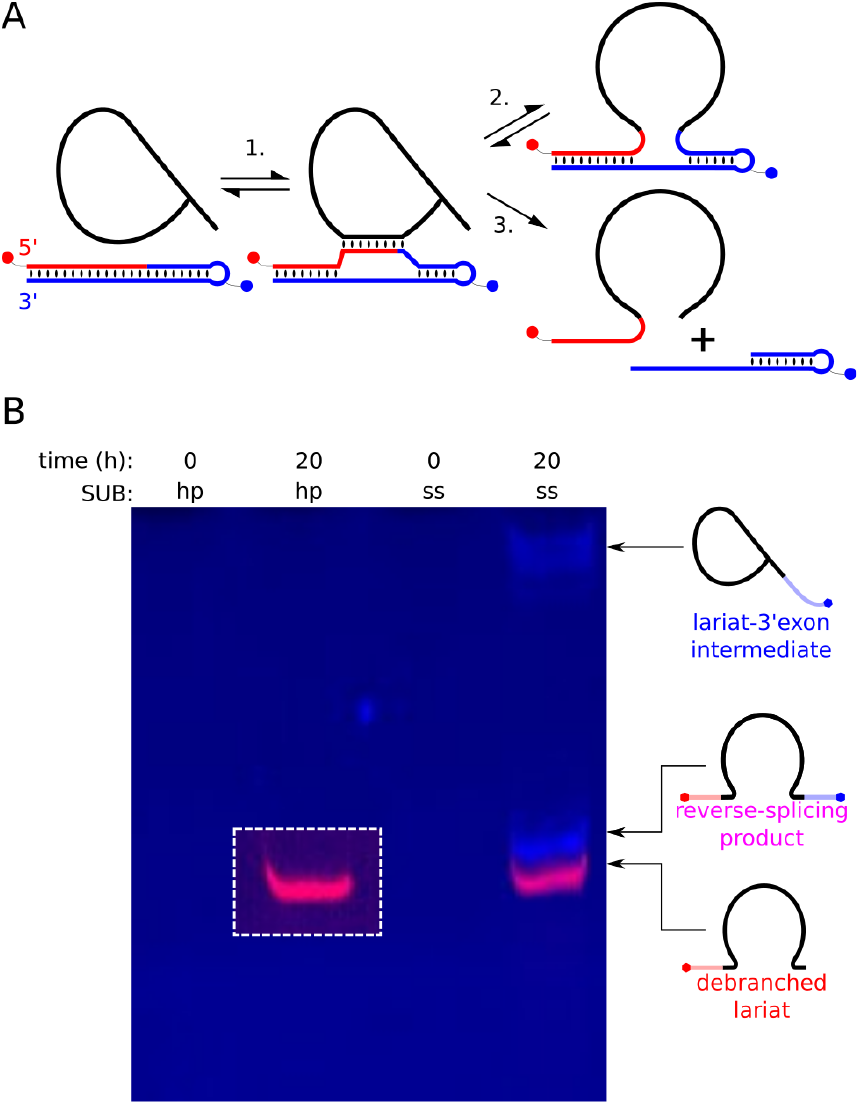
**A)** Illustration of intron lariat reacting with the dsDNA hairpin substrate carrying a 5’-Cy5 label in the 5’-exon (red circle), and an internal FAM modification in the loop of the 3’-exon (blue circle). The lariat unwinds the hairpin DNA at the IBSs (step 1) and catalyzes the reversible reverse-splicing (steps 2 and 3) or quasi-irreversible hydrolysis reaction followed by debranching (steps 4 and 6) **B)** PAGE analysis of reverse splicing into an ssDNA (ss) and the hairpin (hp) dsDNA substrate. The contrast was increased in the region bordered with the white dotted line. Reactions were conducted at 25°C for 20 hours in 200 mM MgCl_2_, 0.5 M NH_4_Cl, 0.001% PEG 8000, and 40 mM Tris•HCl pH 7.5 using 1.5 µM intron lariat and 100 nM of either DNA substrate.

An explanation for the lack of 3’-exon acquisition can be gleaned from the ssDNA time-course (Figure 2) which showed that the debranching product accumulates as hydrolysis progresses in contrast to the transient, short-lived full reverse-splicing product. Additionally, the Mg^2+^ titration experiments revealed that the already low yield and short half-life of the RSP are further reduced at the elevated Mg^2+^ concentrations (Figures 2A and S4) at which the dsDNA experiments were conducted. In conclusion, we surmise that the observed lack of full reverse-splicing product is a result of extensive hydrolysis of the DNA exons because of long incubations at elevated Mg^2+^ concentrations, followed by debranching.

## Discussion

While P.li.LSU.I2 is unlikely to be unique in its capacity for protein-free DNA reverse splicing, it is certainly different in a substantial way from the introns characterized so far that do not show this same capacity. The best biochemically characterized self-splicing group II introns are organellar variants of the IIB1 class (38). The majority of these introns are from a specialized lineage of fungal introns which are either ORF-less or encode a LAGLIDADG homing endonuclease (LHE) (32, 39, 40). This is noteworthy because the ORF-less variants in question are not mobile, while the mobility of the LHE-encoding introns does not involve reverse splicing into DNA (41). In both cases, the introns may experience selection pressure to fold and splice without the help of an encoded maturase, which could explain why they are highly active *in vitro* compared to most other introns. Additionally, it seems plausible that these lineages may have lost the ability to reverse splice into DNA as a result of some evolutionary process. Perhaps it was actively selected against, in order to minimize damage to the host. However, considering that the life cycle of these introns does not involve DNA reverse splicing, mutations that would inactivate this function would be neutral with regard to fitness. As such, it seems more likely that this loss of function was driven by the accumulation of neutral mutations.

Being an organellar variant of the IIB1 class itself, P.li.LSU.I2 is closely related to fungal ORF-less and LHE-encoding introns (28, 41), but unlike them, it is an active RT-encoding retroelement (42, 43), and therefore undoubtedly experiences selection pressure to maintain its capacity to reverse splice into DNA.

The fact that certain protein-free G2Is can reverse splice into, and even hydrolyze DNA warrants their re-evaluation as a biotechnological tool. After the widespread adoption of CRISPR-Cas9, G2I-based tools such as Targetrons (8) have fallen into relative obscurity as genome editing tools. The recent discovery of G2Is described as “hydrolytic endonucleolytic ribozymes” (HYERs) (11), however, suggests that G2Is could be engineered for precise genome editing, offering an alternative to existing technologies with potentially unique advantages. The main current limitation of HYERs is their low sequence recognition capacity of 7 bp and inability to cleave dsDNA (11). The ribozyme we employed in this work overcomes both of these limitations, showing detectable sequence-specific activity with dsDNA, and expanding the recognition sequence to at least 12 nt, though it remains to be seen whether it will retain its activity *in vivo*. Future work may reveal if protein-free activities of the group II intron presented here, including DNA hydrolysis, reverse splicing, and dsDNA unwinding can be improved through directed evolution, thereby offering a versatile alternative to HYERs.

One outstanding question is why P.li.LSU.I2 is capable of protein-free reverse splicing into DNA when all other introns tested thus far are not? Our attempts at identifying the key RNA elements that underpin the ability of some G2Is to reverse splice into DNA did not yield answers. Our initial findings support neither the EBS2a:IBS2a interaction (44), nor the insertion of a single U at position 7 which has the potential to disrupt the ψ-ψ’ interaction (Figure 1A) (15) as factors determining a G2I’s capacity for protein-free debranching with ssDNA (see supplementary results and discussion). By comparing further differences in sequence and structure conservation between groups of G2Is with and without the ability to reverse splice into DNA, it may be possible for future studies to answer this question. Regions of interest that may explain the differences between these introns are the conserved “GACAANGUA” junction between DIC1 and DIC2 (Figure 1B), which is absent in most of the ORF-less and LHE-encoding introns we investigated (19, 32, 39, 41), or the specific stem and loop sequences of DID3(ii), which houses the EBS1 sequence (45, 46).

The origin of the group II intron ribozyme has long been a topic of speculation, with some authors proposing an origin in the RNA world (9, 47–53), while others propose it emerged only after the emergence of the IEP (54). Our data do not support the latter hypothesis, which postulates that G2I IEPs are the primordial core component around which self-splicing ribozymes emerged (54). Rather, our results suggest that the IEP is a late addition to an already fully-fledged self-splicing ribozyme that would have been fully capable of integrating itself into RNA genomes in an RNA world, and surviving the transition into the DNA world without the need for prior association with an RT. It is important to note that this is not at all at odds with the leading “retroelement ancestor” hypothesis for the origin of G2Is (13, 55), which states that all extant G2Is descend from a common ancestor that already encoded an RT, but makes no claims about the origin of the ribozyme or RT prior to the association. We postulate that all the functions of the IEP in splicing are complementary to, and made redundant by pre-existing structures in the ribozyme itself.

This protein takeover was most likely mediated by constructive neutral evolution (CNE) (56, 57). In this scenario, the ancestors of the IEP-dependent ribozymes accumulated inactivating mutations in the regions normally responsible for these functions because the deleterious effects of these mutations were presuppressed by the bound IEP (56, 58). Such an evolutionary course is not without precedent, as CNE is also the leading explanation for the evolution of the spliceosomal machinery from G2Is (58), while similar trends of protein association followed by loss of ribozyme independence can be observed in RNAse P (59, 60) and the ribosomal RNAs (61), both of which can be explained by CNE (56, 57, 62, 63). The most obvious example of this process in group II introns can be seen in the dependence of some introns on the IEP to bind domain VI (DVI) in order to position the branchpoint-A during branching (64). While it seems that this function is ubiquitous among IEPs, most ribozymes also participate in the positioning of DVI by means of a conserved tertiary interaction facilitated by internal loops, bulged nucleotides and wobble pairs in the DVI and DIC1(ii) stems (16, 21, 65, 66). Introns that rely on their IEP for branching usually lack the structures necessary to facilitate these tertiary interactions, and instead have mostly complementary DVI and DIC1(ii) stems (65). The ability to catalyze branching is restored in these introns when the distal stems of DIC1(ii) and DVI are replaced by those of introns capable of branching without the IEP (65).

Another critical role classically attributed to the IEP is the unwinding of dsDNA to allow EBS-IBS pairing (37). Recent work by Monachello et al. has challenged this model, suggesting instead that the ribozyme also participates in duplex unwinding, cooperatively and simultaneously with the IEP (44). Our finding that P.li.LSU.I2 reacts with dsDNA in the absence of its IEP provides the first experimental evidence that the ribozyme is able to unwind its dsDNA target on its own. Thus, even dsDNA duplex unfolding can be interpreted as the IEP merely supporting a pre-existing function of the ribozyme, leaving nothing but reverse transcription as its novel contribution to the RNP. All this suggests that the G2I ribozyme was already fully formed and capable of catalyzing its full repertoire of reactions at the time of its association with the IEP.

The surprising finding that a protein-free group II intron ribozyme is capable of reacting with highly stable DNA duplexes hints at the possibility of early promotor recognition of DNA-dependent RNA polymerase ribozymes. Perhaps ancient polymerase ribozymes functionally resembled *in vitro* evolved RNA polymerase ribozymes that can recognize a promoter sequence (67), with the additional capacity to unwind a double-stranded promoter in a manner similar to the EBS-IBS interaction of group II intron ribozymes.

## Acknowledgements

We would like to thank Alexander Wagner (TU Dortmund University, Germany) for helpful discussions and critical reading of the manuscript, and Jacopo De Capitani (TU Dortmund University, Germany) and Ana Szokoli for helpful discussions.

## References

1. Harris, K.A. and Breaker, R.R. (2018) Large Noncoding RNAs in Bacteria. In Regulating with RNA in Bacteria and Archaea. ASM Press, Washington, DC, USA, pp. 515–526.

2. Peebles, C.L., Perlman, P.S., Mecklenburg, K.L., Petrillo, M.L., Tabor, J.H., Jarrell, K.A. and Cheng, H.-L. (1986) A self-splicing RNA excises an intron lariat. Cell, 44, 213–223.

3. Pyle, A.M. (2016) Group II Intron Self-Splicing. Annu. Rev. Biophys., 45, 183–205.

4. Candales, M.A., Duong, A., Hood, K.S., Li, T., Neufeld, R.A.E., Sun, R., McNeil, B.A., Wu, L., Jarding, A.M. and Zimmerly, S. (2012) Database for bacterial group II introns. Nucleic Acids Res., 40, D187–D190.

5. Zimmerly, S., Guo, H., Eskest, R., Yang, J., Perlman, P.S. and Lambowitz, A.M. (1995) A group II intron RNA is a catalytic component of a DNA endonuclease involved in intron mobility. Cell, 83, 529–538.

6. Yang, J., Zimmerly, S., Perlman, P.S. and Lambowitz, A.M. (1996) Efficient integration of an intron RNA into double-stranded DNA by reverse splicing. Nature, 381, 332–335.

7. Zimmerly, S., Guo, H., Perlman, P.S. and Lambowltz, A.M. (1995) Group II intron mobility occurs by target DNA-primed reverse transcription. Cell, 82, 545–554.

8. Karberg, M., Guo, H., Zhong, J., Coon, R., Perutka, J. and Lambowitz, A.M. (2001) Group II introns as controllable gene targeting vectors for genetic manipulation of bacteria. Nat. Biotechnol., 19, 1162–1167.

9. Lambowitz, A.M. and Belfort, M. (2015) Mobile Bacterial Group II Introns at the Crux of Eukaryotic Evolution. In Mobile DNA III. ASM Press, Washington, DC, USA, pp. 1209–1236.

10. Stamos, J.L., Lentzsch, A.M. and Lambowitz, A.M. (2017) Structure of a Thermostable Group II Intron Reverse Transcriptase with Template-Primer and Its Functional and Evolutionary Implications. Mol. Cell, 68, 926-939.e4.

11. Liu, Z.-X., Zhang, S., Zhu, H.-Z., Chen, Z.-H., Yang, Y., Li, L.-Q., Lei, Y., Liu, Y., Li, D.-Y., Sun, A., et al. (2024) Hydrolytic endonucleolytic ribozyme (HYER) is programmable for sequence-specific DNA cleavage. Science., 383, 1142–1152.

12. King, M.B. and Lapinaite, A. (2024) RNA-based programmable DNA cleavage. Nat. Chem. Biol., 20, 664–665.

13. Toor, N., Hausner, G. and Zimmerly, S. (2001) Coevolution of group II intron RNA structures with their intron-encoded reverse transcriptases. RNA, 7, 1142–52.

14. Wank, H., SanFilippo, J., Singh, R.N., Matsuura, M. and Lambowitz, A.M. (1999) A Reverse Transcriptase/Maturase Promotes Splicing by Binding at Its Own Coding Segment in a Group II Intron RNA. Mol. Cell, 4, 239–250.

15. Haack, D.B., Yan, X., Zhang, C., Hingey, J., Lyumkis, D., Baker, T.S. and Toor, N. (2019) Cryo-EM Structures of a Group II Intron Reverse Splicing into DNA. Cell, 178, 612-623.e12.

16. Xu, L., Liu, T., Chung, K. and Pyle, A.M. (2023) Structural insights into intron catalysis and dynamics during splicing. Nature, 624, 682–688.

17. Matsuura, M. (2001) Mechanism of maturase-promoted group II intron splicing. EMBO J., 20, 7259–7270.

18. Mörl, M. and Schmelzer, C. (1990) Integration of group II intron bl1 into a foreign RNA by reversal of the self-splicing reaction in vitro. Cell, 60, 629–636.

19. Mörl, M., Niemer, I. and Schmelzer, C. (1992) New reactions catalyzed by a group II intron ribozyme with RNA and DNA substrates. Cell, 70, 803–810.

20. Molina-Sánchez, M.D. and Toro, N. (2019) DNA cleavage and reverse splicing of ribonucleoprotein particles reconstituted in vitro with linear RmInt1 RNA. RNA Biol., 16, 930–939.

21. Haack, D.B., Rudolfs, B., Zhang, C., Lyumkis, D. and Toor, N. (2024) Structural basis of branching during RNA splicing. Nat. Struct. Mol. Biol., 31, 179–189.

22. Griffiin, E.A., Qin, Z., Michels, W.J. and Pyle, A.M. (1995) Group II intron ribozymes that cleave DNA and RNA linkages with similar efficiency, and lack contacts with substrate 2’-hydroxyl groups. Chem. Biol., 2, 761–770.

23. Matsuura, M., Saldanha, R., Ma, H., Wank, H., Yang, J., Mohr, G., Cavanagh, S., Dunny, G.M., Belfort, M. and Lambowitz, A.M. (1997) A bacterial group II intron encoding reverse transcriptase, maturase, and DNA endonuclease activities: biochemical demonstration of maturase activity and insertion of new genetic information within the intron. Genes Dev., 11, 2910–2924.

24. Saldanha, R., Chen, B., Wank, H., Matsuura, M., Edwards, J. and Lambowitz, A.M. (1999) RNA and Protein Catalysis in Group II Intron Splicing and Mobility Reactions Using Purified Components. Biochemistry, 38, 9069–9083.

25. Hebbar, S.K., Belcher, S.M. and Perlman, P.S. (1992) A maturase-encoding group MA intron of yeast mitochondria self-splices in vitro. Nucleic Acids Res., 20, 1747–1754.

26. Huang, H.-R., Chao, M.Y., Armstrong, B., Wang, Y., Lambowitz, A.M. and Perlman, P.S. (2003) The DIVa Maturase Binding Site in the Yeast Group II Intron aI2 Is Essential for Intron Homing but Not for In Vivo Splicing. Mol. Cell. Biol., 23, 8809–8819.

27. Fontaine, J.M., Rousvoal, S., Leblanc, C., Kloareg, B. and Loiseaux-de, S.G. (1995) The mitochondrial LSU rDNA of the brown alga Pylaiella littoralis reveals α-proteobacterial features and is split by four group IIB introns with an atypical phylogeny. J. Mol. Biol., 251, 378–389.

28. Costa, M., Fontaine, J.-M., Goër, S.L. and Michel, F. (1997) A group II self-splicing intron from the brown alga Pylaiella littoralis is active at unusually low magnesium concentrations and forms populations of molecules with a uniform conformation. J. Mol. Biol., 274, 353–364.

29. Robart, A.R., Chan, R.T., Peters, J.K., Rajashankar, K.R. and Toor, N. (2014) Crystal structure of a eukaryotic group II intron lariat. Nature, 514, 193–197.

30. Zuker, M. (2003) Mfold web server for nucleic acid folding and hybridization prediction. Nucleic Acids Res., 31, 3406–3415.

31. Lucas, J.K., Gruenke, P.R. and Burke, D.H. (2023) Minimizing amplification bias during reverse transcription for in vitro selections. RNA, 29, 1301–1315.

32. Liu, T. and Pyle, A.M. (2021) Discovery of highly reactive self-splicing group II introns within the mitochondrial genomes of human pathogenic fungi. Nucleic Acids Res., 49, 12422–12432.

33. Moran, J. V, Zimmerly, † Steven, Eskes, R., Kennell, J.C., Lambowitz, A.M., Butow, R.A. and Perlman, P.S. (1995) Mobile group II introns of yeast mitochondrial DNA are novel site-specific retroelements. Mol. Cell. Biol., 15, 2828–2838.

34. Jarrell, K.A., Peebles, C.L., Dietrich, R.C., Romiti, S.L. and Perlman, P.S. (1988) Group II intron self-splicing. Alternative reaction conditions yield novel products. J. Biol. Chem., 263, 3432–3439.

35. Roitzsch, M. and Pyle, A.M. (2009) The linear form of a group II intron catalyzes efficient autocatalytic reverse splicing, establishing a potential for mobility. RNA, 15, 473–482.

36. Aizawa, Y., Xiang, Q., Lambowitz, A.M. and Pyle, A.M. (2003) The Pathway for DNA Recognition and RNA Integration by a Group II Intron Retrotransposon. Mol. Cell, 11, 795–805.

37. Singh, N.N. and Lambowitz, A.M. (2001) Interaction of a group II intron ribonucleoprotein endonuclease with its DNA target site investigated by DNA footprinting and modification interference. J. Mol. Biol., 309, 361–386.

38. Sigel, R.K.O. (2005) Group II intron ribozymes and metal ions - A delicate relationship. Eur. J. Inorg. Chem., 2005, 2281–2292.

39. Mullineux, S.-T., Costa, M., Bassi, G.S., Michel, F. and Hausner, G. (2010) A group II intron encodes a functional LAGLIDADG homing endonuclease and self-splices under moderate temperature and ionic conditions. RNA, 16, 1818–1831.

40. van der Veen, R., Arnberg, A.C., van der Horst, G., Bonen, L., Tabak, H.F. and Grivell, L.A. (1986) Excised group II introns in yeast mitochondria are lariats and can be formed by self-splicing in vitro. Cell, 44, 225–234.

41. Toor, N. and Zimmerly, S. (2002) Identification of a family of group II introns encoding LAGLIDADG ORFs typical of group I introns. RNA, 8, 1373–1377.

42. Ikuta, K., Kawai, H., Müller, D.G. and Ohama, T. (2008) Recurrent invasion of mitochondrial group II introns in specimens of Pylaiella littoralis (brown alga), collected worldwide. Curr. Genet., 53, 207–216.

43. Zerbato, M., Holic, N., Moniot-Frin, S., Ingrao, D., Galy, A. and Perea, J. (2013) The Brown Algae Pl.LSU/2 Group II Intron-Encoded Protein Has Functional Reverse Transcriptase and Maturase Activities. PLoS One, 8, e58263.

44. Monachello, D., Lauraine, M., Gillot, S., Michel, F. and Costa, M. (2021) A new RNA–DNA interaction required for integration of group II intron retrotransposons into DNA targets. Nucleic Acids Res., 49, 12394–12410.

45. Skilandat, M. and Sigel, R.K.O. (2014) The Role of Mg(II) in DNA Cleavage Site Recognition in Group II Intron Ribozymes. J. Biol. Chem., 289, 20650–20663.

46. Steffen, F.D., Khier, M., Kowerko, D., Cunha, R.A., Börner, R. and Sigel, R.K.O. (2020) Metal ions and sugar puckering balance single-molecule kinetic heterogeneity in RNA and DNA tertiary contacts. Nat. Commun., 11, 104.

47. Gilbert, W. (1986) Origin of life: The RNA world. Nature, 319, 618.

48. Lambowitz, A.M. and Perlman, P.S. (1990) Involvement of aminoacyl-tRNA synthetases and other proteins in group I and group II intron splicing. Trends Biochem. Sci., 15, 440–444.

49. Dai, L. and Zimmerly, S. (2003) ORF-less and reverse-transcriptase-encoding group II introns in archaebacteria, with a pattern of homing into related group II intron ORFs. RNA, 9, 14–19.

50. Fine, J.L. and Pearlman, R.E. (2023) On the origin of life: an RNA-focused synthesis and narrative. RNA, 29, 1085–1098.

51. Zhao, C. and Pyle, A.M. (2016) Crystal structures of a group II intron maturase reveal a missing link in spliceosome evolution. Nat. Struct. Mol. Biol., 23, 558–565.

52. McNeil, B.A., Semper, C. and Zimmerly, S. (2016) Group II introns: versatile ribozymes and retroelements. WIREs RNA, 7, 341–355.

53. Koonin, E. V, Senkevich, T.G. and Dolja, V. V (2006) The ancient Virus World and evolution of cells. Biol. Direct, 1, 29.

54. Curcio, M.J. and Belfort, M. (1996) Retrohoming: cDNA-Mediated Mobility of Group II Introns Requires a Catalytic RNA. Cell, 84, 9–12.

55. Zimmerly, S. and Semper, C. (2015) Evolution of group II introns. Mob. DNA, 6, 7.

56. Stoltzfus, A. (1999) On the Possibility of Constructive Neutral Evolution. J. Mol. Evol., 49, 169–181.

57. Muñoz-Gómez, S.A., Bilolikar, G., Wideman, J.G. and Geiler-Samerotte, K. (2021) Constructive Neutral Evolution 20 Years Later. J. Mol. Evol., 89, 172–182.

58. Koonin, E. V. (2016) Splendor and misery of adaptation, or the importance of neutral null for understanding evolution. BMC Biol., 14, 114.

59. Guerrier-Takada, C., Gardiner, K., Marsh, T., Pace, N. and Altman, S. (1983) The RNA moiety of ribonuclease P is the catalytic subunit of the enzyme. Cell, 35, 849–857.

60. Evans, D., Marquez, S.M. and Pace, N.R. (2006) RNase P: interface of the RNA and protein worlds. Trends Biochem. Sci., 31, 333–341.

61. Xu, D. and Wang, Y. (2021) Protein-free ribosomal RNA scaffolds can assemble poly-lysine oligos from charged tRNA fragments. Biochem. Biophys. Res. Commun., 544, 81–85.

62. Schencking, I., Rossmanith, W. and Hartmann, R.K. (2020) Diversity and Evolution of RNase P. In Evolutionary Biology—A Transdisciplinary Approach. Springer International Publishing, Cham, pp. 255–299.

63. Lukeš, J., Archibald, J.M., Keeling, P.J., Doolittle, W.F. and Gray, M.W. (2011) How a neutral evolutionary ratchet can build cellular complexity. IUBMB Life, 63, 528–537.

64. Smathers, C.M. and Robart, A.R. (2020) Transitions between the steps of forward and reverse splicing of group IIC introns. RNA, 26, 664–673.

65. Monachello, D., Michel, F. and Costa, M. (2016) Activating the branch-forming splicing pathway by reengineering the ribozyme component of a natural group II intron. RNA, 22, 443–455.

66. Li, C.-F., Costa, M. and Michel, F. (2011) Linking the branchpoint helix to a newly found receptor allows lariat formation by a group II intron. EMBO J., 30, 3040–3051.

67. Cojocaru, R. and Unrau, P.J. (2021) Processive RNA polymerization and promoter recognition in an RNA World. Science., 371, 1225–1232.

