## Supplementary Information for "Protein-Free Catalysis of DNA Hydrolysis and Self-Integration by a Ribozyme"

### Supplementary methods

#### Isolated DNA debranching time-course

Prior to initiating the reaction, 100 nM of intron lariat was denatured in 40 mM Tris•HCl pH 7.5 and 0.1% Tween 20 (Sigma-Aldrich) by heating to 90°C for 1 minute, then slow cooling (0.1°C/s) to 25°C. The intron was then re-folded by addition of 0.5 M $NH_4Cl$ and 50 mM $MgCl_2$ and incubating at 25°C for 10 minutes. Tubes containing the refolded intron were transferred to an ice bath before adding 1 µM of 5'-Cy5 labeled 5'-exon DNA "DNA dbr sub." (Table S1), and a sample was taken for the t=0 timepoint. To start the reaction the tubes were transferred into a 25°C thermocycler, and further timepoints were taken over the course of the reaction. Samples were quenched in four volumes of denaturing formamide gel-loading buffer (96% formamide, 20 mM EDTA pH 8, bromophenol blue) and kept on ice. Samples were electrophoresed on a single-layer 6% gel as described in the main text (See materials and methods subsection "Denaturing polyacrylamide gel electrophoresis").

#### Reverse-splicing time-course at 350 mM $MgCl_2$

The reverse-splicing time-course at 350 mM $MgCl_2$ was performed in an identical way to the 50 mM $MgCl_2$ reverse-splicing time-course with 200 nM DNA substrate that was described in the main text (See materials and methods subsection "Reverse-splicing reactions"), except that 3.2 µl samples were quenched in 30.6 µl of quench buffer (92% formamide, 38.82 mM EDTA pH 8, bromophenol blue).

#### Time-course data analysis

The isolated debranching time-course displayed a biphasic kinetic behavior and was fitted with a biphasic decay:

$$[D] = [D]_\infty \cdot F_{fast} \cdot (1 - e^{-k_{fast} \cdot t}) + [D]_\infty \cdot F_{slow} \cdot (1 - e^{-k_{slow} \cdot t})$$

(Eq. S1)

Where $[D]$ is the molar concentration of the debranched product at any timepoint $t$, $[D]_\infty$ is the value of $[D]$ as $t$ approaches infinity, $F_{fast}$ is the fraction of the fast-reacting population (value between 0 and 1), $F_{slow}$ is the fraction of the slow-reacting population, where $F_{slow} = 1 - F_{fast}$, and $k_{fast}$ and $k_{slow}$ are the apparent rate constants of the fast and slow reaction phase, respectively.

The data from the reverse-splicing time-course at 350 mM $MgCl_2$ was fitted with the same equation as the 50 mM $MgCl_2$ time-course data (main text Eq. 4).

### Supplementary results and discussion

#### Key factors responsible for protein-free group II intron reverse splicing into DNA remain elusive

It may be possible for future studies to determine the key RNA elements that underpin the ability of some G2Is to reverse splice into DNA by comparing their sequence and structure conservation with that of G2Is that lack this ability. For example, RT-encoding class IIB G2Is contain a highly conserved nucleotide “EBS2a” that forms a base pair with their DNA targets named “IBS2a” (Figure S6C) (1). There is ample evidence that EBS2 is mostly involved in retrohoming, which is additionally supported by the loss of EBS2 in many ORF-less and LHE-encoding class CL1 introns (1, 2). Inspecting the sequences of the ORF-less and LHE-encoding class CL1 introns from the mitochondria of fungi, which were presumed unable to reverse splice into DNA, confirmed that they lacked the capacity to form the EBS2a:IBS2a interaction, with the exception of the *S. cerevisiae* intron “bI1”. Investigating the resolved structures of CL1 introns, we also noticed that they contain a conserved U:G wobble pair in subdomain ID(iv), which supports the EBS2a:IBS2a interaction. With the exception of bI1, this wobble pair was absent in the inspected ORF-less and LHE-encoding introns. Interestingly, however, we noted that in the original 1992 study on bI1 by Mörl *et al*, in which bI1 failed to reverse splice into DNA, the substrate used was missing the IBS2a nucleotide (3), which we hypothesized may have been the reason for the author’s negative findings.

Reverse-splicing experiments in which the group II intron bI1 was reacted with DNA substrates were in agreement with previous reports, as partial, but not complete reverse splicing were achieved with a DNA substrate which was designed to restore the previously missing EBS2a:IBS2a interaction (Figure S7). However, isolated debranching experiments with 5’-exons designed to restore the EBS2a:IBS2a interaction produced a minuscule amount of product, which was not detected with the original sequence lacking the interaction (Figure S7). In contrast, isolated debranching experiments with P.li.LSU.I2 where the location and presence of the IBS2a nucleotide were varied showed no discernable effect on debranching (Figure S6D). These results suggest that an as-of-yet uncharacterized sequence component of the substrate, which is distinct from that of the EBS2a:IBS2a interaction, may have an effect on reverse splicing.

Structural studies have identified a conserved tertiary contact (ψ-ψ’) between A341 (ψA) and a U:G wobble pair in Domain V that is required for branching. We hypothesized that the uracil inserted opposite of the ψA in bI1 may prevent the re-establishment of the ψ-ψ’ interaction by base-pairing with the ψA after forward-branching. This hypothesis was disproven when we inserted a uridine residue at the cognate site of P.li.LSU.I2 (Figure S6A), and found that it did not affect reverse splicing (Figure S6B). As such, we do not currently have a mechanistic explanation for what makes a G2I capable of fully reverse splicing into DNA in the absence of an IEP. Regions of interest that may explain the differences between these introns are the conserved “GACAANGUA” junction between DIC1 and DIC2 (which is absent in most of the ORF-less and LHE-encoding introns we investigated) (3–6), or the specific stem and loop sequences of DID3(ii), which houses the EBS1 sequence (7, 8).

### Supplementary figures:

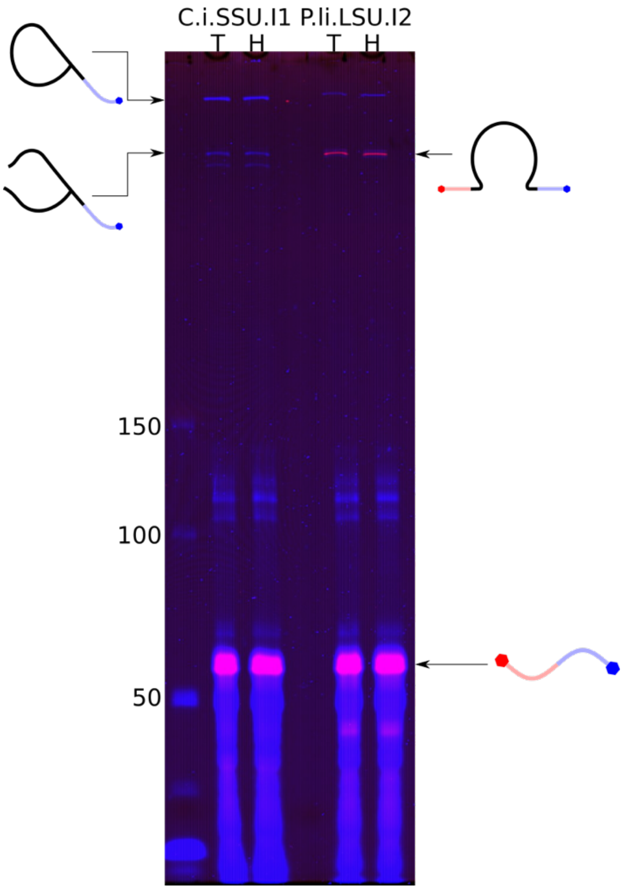

***Figure S1: Comparison of reverse-splicing reactions between P.li.LSU.I2 and C.i.SSU.I1.*** *Both introns were incubated under identical reaction conditions, in either a Tris (T) or HEPES (H) buffer (50 mM of either Tris•HCl pH 7.5 or HEPES•KOH pH 7.5, 0.5 µM lariat, 100 mM KCl, 200 mM $MgCl_2$, and 5 µM DNA substrate "DNA RS sub. 2") for 22 h at 25°C. While both introns can acquire the 3'-exon, only P.li.LSU.I2 is capable of debranching with the DNA 5'-exon. The two bands of a similar MW to the RSP seen in the C.i.SSU.I1 reactions that only carry the 3'-exon FAM fluorophore are most likely the broken lariat-3'-exon intermediate and the product of a linearized intron performing reverse-ligation.*

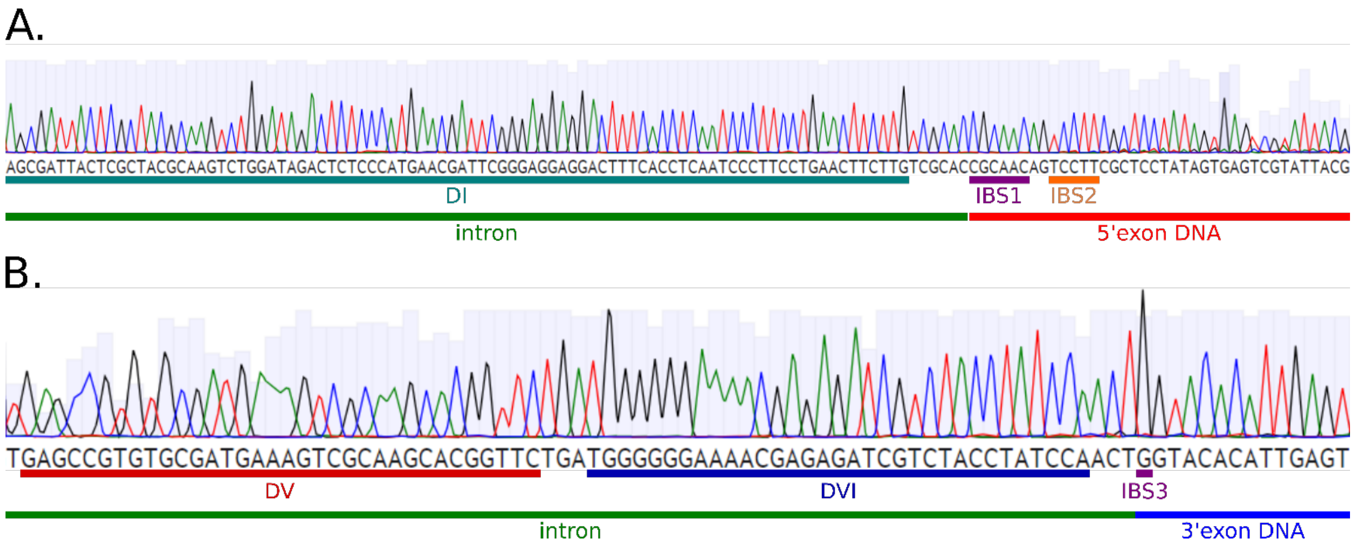

***Figure S2: Sequencing results of "DNA RS sub. 2" ssDNA reverse-splicing RT-PCR. A)*** *The reverse complement of the 5'-splice-site. The intron is sequence-specifically inserted directly downstream of IBS1.* ***B)*** *The intron is sequence-specifically inserted at the 3'-splice-site, directly upstream of IBS3.*

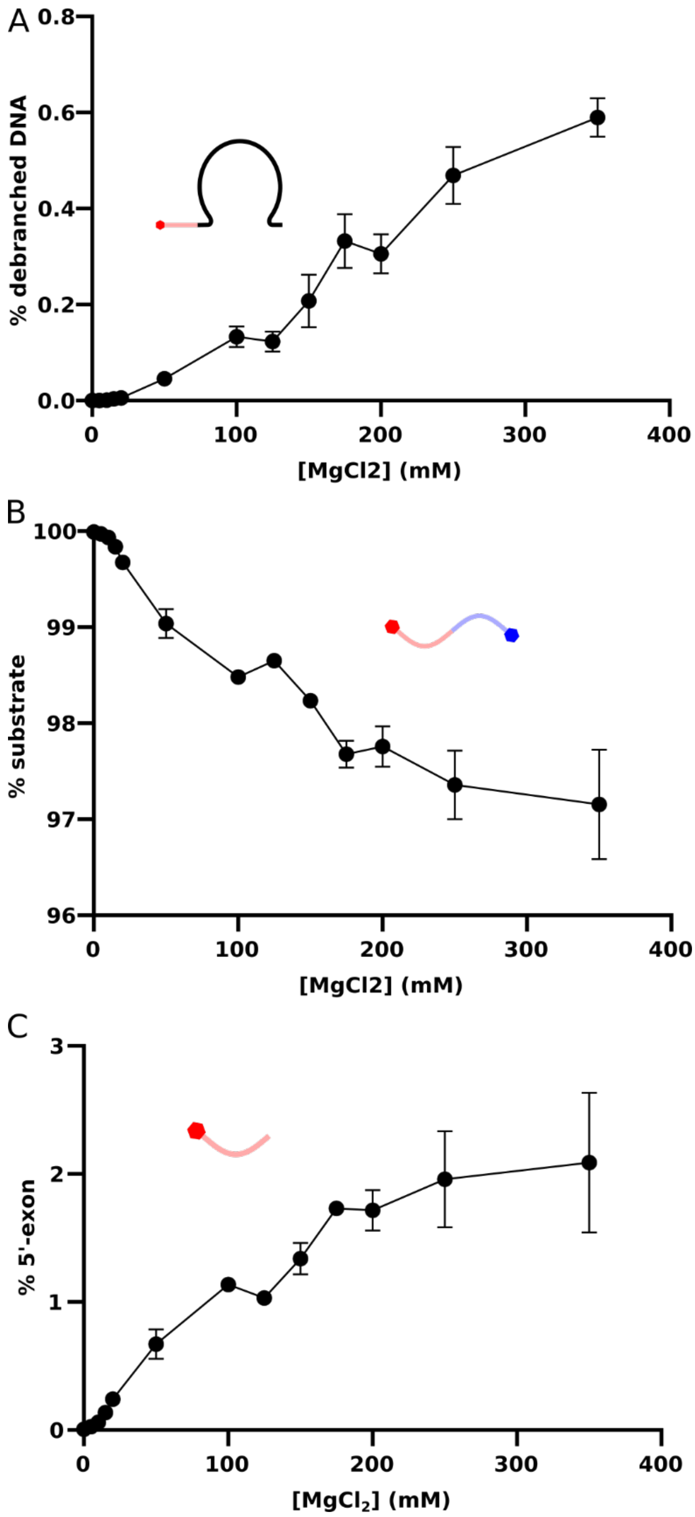

***Figure S3: Molecular species other than the RSP from the $MgCl_2$ titration experiment in Figure 2A. A)*** *Debranched product.* ***B)*** *DNA substrate.* ***C)*** *5’-exon DNA.*

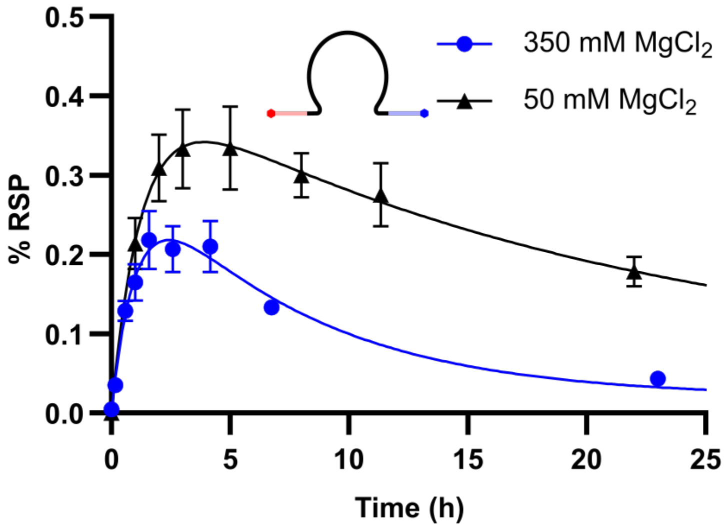

***Figure S4: Comparison of reverse-splicing product (RSP) time courses at 50 mM and 350 mM $MgCl_2$.*** *Experiments were conducted in a buffer containing 40 mM Tris•HCl pH 7.5, 0.001% PEG 8000, 0.5 M $NH_4Cl$, 100 nM intron lariat and 200 nM DNA substrate, and the specified concentration of $MgCl_2$. Datapoints are the averages of triplicates. Error bars represent ± standard deviations (n = 3).*

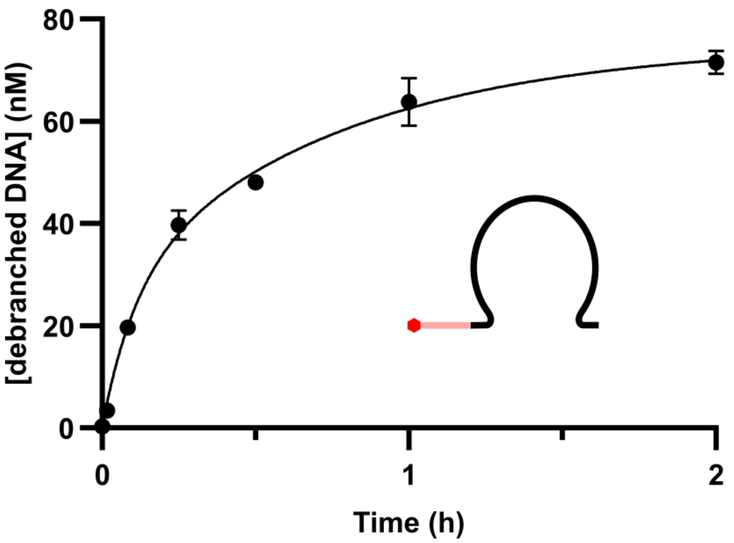

***Figure S5: Debranching time course of P.li.LSU.I2 lariat with a 5'-exon DNA substrate.*** *The reactions were incubated at 25°C and contained 40 mM Tris•HCl pH 7.5, 0.1% Tween 20, 0.5 M $NH_4Cl$, 50 mM $MgCl_2$, 100 nM intron lariat and 1 µM 5'-Cy5 labeled DNA exon. Datapoints are the averages of triplicates. Error bars represent ± standard deviations (n = 3).*

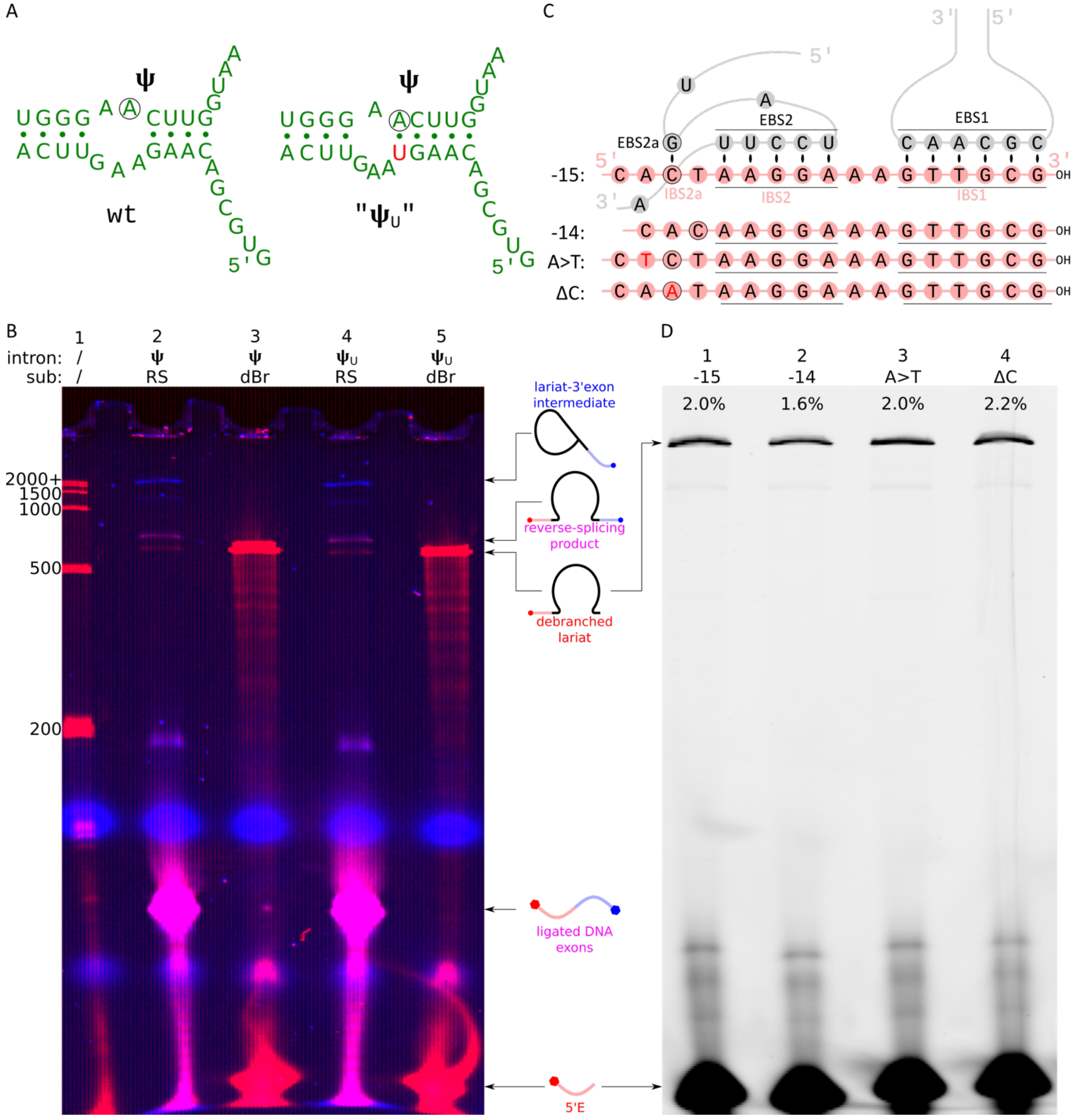

***Figure S6: Mutational studies of protein-free reverse splicing into DNA.*** ***A)*** *Schematic of a section P.li.LSU.I2. On the left is the wild-type sequence. On the right is the "$\psi_U$" mutant, in which the inserted U on the bottom strand has the potential to base pair with the ψA, potentially disabling the ψ- ψ' tertiary interaction.* ***B)*** *Denaturing PAGE of the wild type and "$\psi_U$" mutants debranching (dBr) and reverse splicing (RS). Reactions were carried out in the standard protocol described in the main text. Reactions contained 40 mM Tris•HCl pH 7.5, 0.001% PEG 8000, 0.5 M $NH_4Cl$, 100 mM $MgCl_2$ 100 nM intron lariat and 200 nM DNA substrate or 5'-exon DNA, and were incubated at 25°C for 30 minutes.* ***C)*** *Top: Schematic of the EBS2:IBS2 and EBS2a:IBS2a interaction between the modified P.li.LSU.I2 used in this study (gray), and the standard 5'-exon DNA substrate (red) in which the IBS2a nucleotide (black outline) is in the -15 position (15 bases upstream of the insertion site). Bottom: 5'-exon variants used to study the effect of the EBS2a:IBS2a interaction. Substrate "-14" has a point deletion, moving IBS2a directly upstream of IBS2. Substrate "A>T" has the A at position -16 (which is a conserved purine thought to interact with a conserved amino acid in the IEP, and be involved in retrohoming* (1)*) mutated to a T. Substrate "ΔC" has the IBS2a base (in this case a C) mutated to an A, abolishing the EBS2a:IBS2a interaction.* ***D)*** *Denaturing PAGE of debranching reactions with the four substrate variants described in C). Percentages represent the percent conversion of DNA substrate to the debranching product after 20 minutes at 25°C (40 mM*

*Tris•HCl pH 7.5, 0.1% Tween 20, 0.5 M $NH_4Cl$, 50 mM $MgCl_2$, 0.5 µM intron lariat and 5 µM 5′-6FAM labeled DNA substrate).*

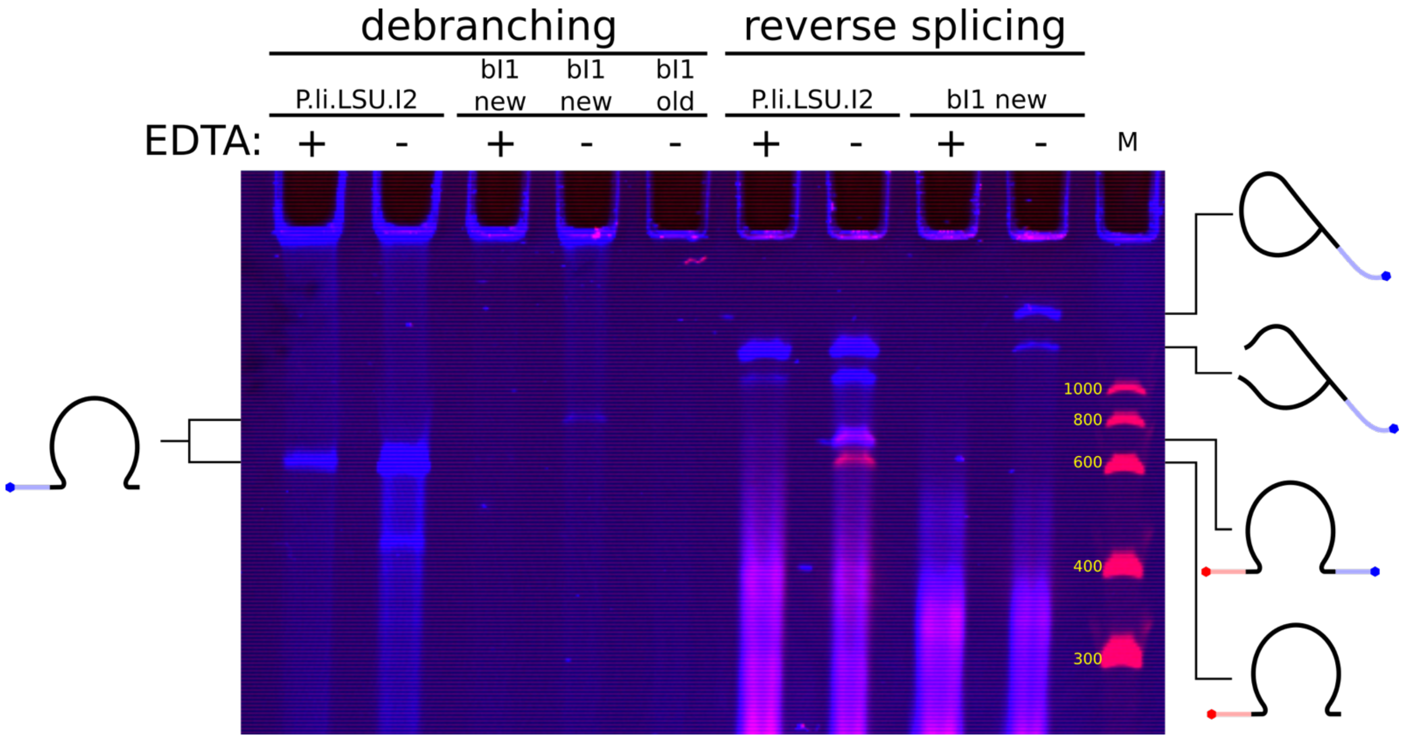

***Figure S7: Debranching and reverse splicing of P.li.LSU.I2 and bI1.*** *Both reactions were carried out under identical conditions (30 mM Tris•HCl pH 7.5, 0.001% PEG 8000, 100 mM MgCl$_2$, 1 M NH$_4$Cl, 250 nM lariat, 2.5 µM DNA substrate). Negative controls were mixed with 125 mM EDTA before the addition of substrate. All samples were incubated for 1 h at 25°C. The debranching substrate was 5'-6FAM labeled, hence the blue color. The bI1 intron was reacted with either the 5'-exon sequence used in Mörl et al (3), ("bI1 old"), or a new design that minimizes secondary structures ("bi1 new"). The leftmost lane contains a Cy5 labeled Riboruler low-range RNA ladder (Thermo). A faint band corresponding to the expected size of the debranched product can be seen for the "bi1 new" debranching substrate. Complete reverse splicing is not detected for bI1 with the new substrate, but we were able to detect the lariat-3'-exon intermediate, as well as the broken intermediate, which is in agreement with the results of Mörl et al.*

### Table S1: nucleic acid sequences

| **A) IVT Templates:** | |
|---|---|
| mod. P.li.LSU.I2 IVT template[a] | GCG*TAATACGACTCACTATAG*GAGCGAAGGACTGTTGCG**GTGCGACAAGAAGT**<br>**TCAGGAAGGGATTGAGGTGAAAGTCCTCCCGAATCGTTCATGGGAGAGTC**<br>**TATCCAGACTTGCGTAGCGAGTAATCGCTAGGTGAGAAGCTCTGGAGACAATG**<br>**TACCTGCCCTTCAATTGGAGGTGCCAGGGCTAGGCTTTGTTCCTATGGCTAGCG**<br>**AAAGCGAATACAGGCTATTAGGCGTCGTGATCCTTAAACTCTTTCGACATAAGG**<br>**GAGAGAAAAGTAGATGAGTTACCCTCCCGCAACTGGGGTAACTGAACGATCAA**<br>**TATCTACCGGCGTACACTGTGAGCCTAAAGGAAACGAATGAAAGTGTCCCTTCC**<br>**TGGGAACTTGGTAAGCCCAATAGACTCCCTCTGGGAAACCAGAGGAGGAGTA**<br>**AGAGCAATCTTACTTATCGTCTAGTGGGTAAAAGACATAATAGGAAGCAAATG**<br>**CTTGGTTGTAACGATCGAGATAGAATCTGTTGATTTAAGCTGAAAGGCTGCAG**<br>**ACTTATTAAATGGGTGTTCTGCTTTCGGCAGCAGAATGCGTGAGCCGTGTGCGA**<br>**TGAAAGTCGCAAGCACGGTTCTGATGGGGGGAAAACGAGAGATCGTCTACCT**<br>**ATCCAACT**GCGTTCACTGAGGGTCCTAAGATT |
| mod. P.li.LSU.I2 $\psi_U$ IVT template[a,b] | GCGAAAT*TAATACGACTCACTATAG*GGAAATACTAAGGACTGTTGCG**GTGCGAC**<br>**AAGTAAGTTCAGGAAGGGATTGAGGTGAAAGTCCTCCCGAATCGTTCATG**<br>**GGAGAGTCTATCCAGACTTGCGTAGCGAGTAATCGCTAGGTGAGAAGCTCTGG**<br>**AGACAATGTACCTGCCCTTCAATTGGAGGTGCCAGGGCTAGGCTTTGTTCCTAT**<br>**GGCTAGCGAAAGCGAATACAGGCTATTAGGCGTCGTGATCCTTAAACTCTTTCG**<br>**ACATAAGGGAGAGAAAAGTAGATGAGTTACCCTCCCGCAACTGGGGTAACTG**<br>**AACGATCAATATCTACCGGCGTACACTGTGAGCCTAAAGGAAACGAATGAAAG**<br>**TGTCCCTTCCTGGGAACTTGGTAAGCCCAATAGACTCCCTCTGGGAAACCAGAG**<br>**GAGGAGTAAGAGCAATCTTACTTATCGTCTAGTGGGTAAAAGACATAATAGGA**<br>**AGCAAATGCTTGGTTGTAACGATCGAGATAGAATCTGTTGATTTAAGCTGAAA**<br>**GGCTGCAGACTTATTAAATGGGTGTTCTGCTTTCGGCAGCAGAATGCGTGAGC**<br>**CGTGTGCGATGAAAGTCGCAAGCACGGTTCTGATGGGGGGAAAACGAGAGAT**<br>**CGTCTACCTATCCAACT**GCCGGTGCGCATAACCACCTCAGTGCGAGCAAGGAAA<br>TCATGGTTTTTACCTCCTGAATTCGGATCCCTCGAGCGATACACACTTCTATAGTG<br>TCACCTAAATGCGTTTAAACCTTCCTGCAGGTGACGATTACCTAACAATCGGTCG<br>ATTCGTTTGATGTTATGTTTTGTTCTCGCTTTGGTTGGCAGGT |
| C.i.SSU.I1 IVT template | GCG*TAATACGACTCACTATAG*GAGCGAAGGAT**GTGCGACTTGTTAAGTTTTAAC**<br>**AAAAATTGTATAACGTTTATTAATGATTATACATTGTATTTCATCTTACAATAGC**<br>**CTAATTAGATATGCATTTAGGGTAACTTTTTTGTATAAAGCTCTAATTATAAGTG**<br>**TAAATACACTTTTAGGCTTCTTCTATGGTTAGAGAAATCGAACCAATGTAATTA**<br>**AAACTTTGATGTATTAGGCATTTAACGTGTCCTTGGTTAAATGAAGATGAACAT**<br>**AAGTATACAAAGTAAAATTGGAACCTAAGGAAGAATTGTTTTTGTTAAGAAAC**<br>**AAGGTAATACCTATAACTGGCTAATATAAATTTGCAAGGTTTATTGTAAAATAA**<br>**ACTATAGGTTAGAGGTAAAAGGATAATGTAAAAAGCGAATGCAATTCTGTAAT**<br>**GGAATTGATAGGGTATATACCTAACTTGAAAGAGTGCTGACTTACATATAGAT**<br>**GTTATTTACGTTTCGACGTAAATAATGTTTGAGCCGTATGCTATGAAAGTAGCA**<br>**CGTACGGTTCTAAGAGGGGGAAAGTCCGAGAGGACCTACCTATCTCAAC**TGCG<br>TTCACTGAGGGTCCTAAGATT |
| bI1 IVT template | GCG*TAATACGACTCACTATAG*GGCGAATTGGGCCCGACGTCGCATGCTCCCGGC<br>CGCCATGGCGGCCGCGGGAATTCGATTTGTTTATGGACAGA**GTGAGACAAGTAT**<br>**AAGTATATTATTATAATATCATACCATTAAATAAATTATTTTAATGAAATGATTA**<br>**TGTTTATATATAACATATACCTAATTAGACATGCATTATTAGTAATAATTTTGTA**<br>**TGAAACTCTAATAATAATAATTATTATTAATTATTAAGGTAAGATTCATATgGAT**<br>**AGCGTAAGTCAATCTAATATTATAAAATATCGTAACATAAACAATATTTTTTTCT**<br>**ATTATTAATTAATAAATAATAATAAATAAAAATAATTATATGAGAAGTAAGAT**<br>**ATTCAATTCTGTCTAGAATACATATATATACGTTAATACTCATCGGTATAAAATT**<br>**AGAATCCTAAGTGAATTATTGAAAGTATAATAATATAAACTTGGTAAGCCCAA**<br>**TTATTTCCATATAATATTAATATAAATATTATATGGTAGTTATATATAATATTAT** |

| | |
|---|---|
| | **TAAATAAATAATAATAGAAATTATAATATAGATAAGTGGGTAAAAGACTATTGAAAAAGCTAAAGATTATATGTAATGTATAATATAGATCAAATTATTTATATATTTTAATAAAAATATATTAATAATGGTTAATATTATTATTAATTAATTAATTAATTAATTAATAATAATAACGAATAAATGATTAATGTGAAAGCATGCTAACTTCAATATAGGATGATTTATATAGTATATAAATTGTTTGAGCTGTATACTATGAAAGTAGTACGTACAGTTCTGAGTGGGGGAAAATTTGTAAAGATCTACCTATCACAAT**TGTCACATTGAATCACTAGTGAATTCGCGGCCGCCTGCAGGTCGACCATATGGGAGAGCTCCCAACGCGTTGGATGCATAGCTTGAGTATTCTATAGTGTCACCTAAATAGCTTGGCGTAATCATGGTCATAGCTGTTTCCTGTGTGAAATTGTTATCCGCT |
| **B) DNA Substrates:** | |
| DNA RS sub. 1[c,d] | /5Cy5/AAAAACGAAGGACTGTTGCGGAAATAAATAAATAAATAAATAAATAAATAAATAAATAAATAAATAAATAAATAAATAAATAAATAAATAAATAAA/36-FAM/ |
| DNA RS sub. 2[c,d] | GTGAGTCG/iCy5/GCGTAATACGACTCACTATAGGAGCGAAGGACTGTTGCGGGTACACATTGAGT/36-FAM/ |
| DNA RS hp sub. [c,d] | /5Cy5/CGGGAGATCTGGACGAAGGAAAGTTGCGGATACACTGAGTGACG/iFluorT/AGTCACTCAGTGTATCCGCAACTTTCCTTCGTCCAGATCTCCCG |
| DNA dbr sub.[c,d] | /5Cy5/CCATTAGAGCGTAAGCTCATCACTAAGGAAAGTTGCG |
| DNA dbr sub. -15[c,d] | /56-FAM/CCATTAGAGCGTAAGCTCATCACTAAGGAAAGTTGCG |
| DNA dbr sub. -14[c,d] | /56-FAM/CCATTAGAGCGTAAGCTCATCACAAGGAAAGTTGCG |
| DNA dbr sub. A>T[c,d] | /56-FAM/CCATTAGAGCGTAAGCTCATCTCTAAGGAAAGTTGCG |
| DNA dbr sub. ΔC[c,d] | /56-FAM/CCATTAGAGCGTAAGCTCATCAATAAGGAAAGTTGCG |
| Plasmid pS (BspHI cut)[d,e] | CATGAGCGGATACATATTTGAATGTATTTAGAAAAATAAACAAATAGGGGTTCCGCGCACATTTCCCCGAAAAGTGCCACCTGATGCGGTGTGAAATACCGCACAGATCGATCCCGCGAAATTAATACGACTCACTATAGGGGAATTGTGAGCGGATAACAATTCCCCTCTAGAAATAATTTTGTTTAACTTTAAGAAGGAGATATACCAAAGGCCAGCAAAAGGCCAGGAACCGTAAAAAGGCCGCGTTGCTGGCGTTTTTCCATAGGCTCCGCCCCCCTGACGAGCATCACAAAAATCGACGCTCAAGTCAGAGGTGGCGAAACCCGACAGGACTATAAAGATACCAGGCGTTTCCCCCTGGAAGCTCCCTCGTGCGCTCTCCTGTTCCGACCCTGCCGCTTACCGGATACCTGTCCGCCTTTCTCCCTTCGGGAAGCGTGGCGCTTTCTCATAGCTCACGCTGTAGGTATCTCAGTTCGGTGTAGGTCGTTCGCTCCAAGCTGGGCTGTGTGCACGAACCCCCCGTTCAGCCCGACCGCTGCGCCTTATCCGGTAACTATCGTCTTGAGTCCAACCCGGTAAGACACGACTTATCGCCACTGGCAGCAGCCACTGGTAACAGGATTAGCAGAGCGAGGTATGTAGGCGGTGCTACAGAGTTCTTGAAGTGGTGGCCTAACTACGGCTACACTAGAAGAACAGTATTTGGTATCTGCGCTCTGCTGAAGCCAGTTACCTTCGGAAAAAGAGTTGGTAGCTCTTGATCCGGCAAACAAACCACCGCTGGTAGCGGTGGTTTTTTTGTTTGCAAGCAGCAGATTACGCGCAGAAAAAAAGGATCTCAAGAAGATCCTTTGATCTTTTCTACGGGGTCTGACGCTCAGTGGAACGAAAACTCACGTTAAGGGATTT**ATCGGGAGATCTGGACGAAGGAAAGTTGCGGATACACATTGAGTGATTTC**GAGTAAACTTGGTCTGACAGTTACCAATGCTTAATCAGTGAGGCACCTATCTCAGCGATCTGTCTATTTCGTTCATCCATAGTTGCCTGACTCCCCGTCGTGTAGATAACTACGATACGGGAGGGCTTACCATCTGGCCCCAGTGCTGCAATGATACCGCGAGACCCACGCTCACCGGCTCCAGATTTATCAGCAATAAACCAGCCAGCCGGAAGGGCCGAGCGCAGAAGTGGTCCTGCAACTTTATCCGCCTCCATCCAGTCTATTAATTGTTGCCGGGAAGCTAGAGTAAGTAGTTCGCCAGTTAATAGTTTGCGCAACGTTGTTGCCATTGCTACAGGCATCGTGGTGTCACGCTCGTCGTTTGGTATGGCTTCATTCAGCTCCGGTTCCCAACGATCAAGGCGAGTTACATGATCCCCCATGTTGTGCAAAAAAGCGGTTAGCTCCTTCGGTCCTCCGATCGTTGTCAGAAGTAAGTTGGCCGCAGTGTTATCACTCATGGTTATGGCAGCACTGCATAATTCTCTTACTGTCATGCCATCCGTAAGATGCTTTTCTGTGACTGGTGAGTACTCAACCAAGTCATTCTGAGAATAGTGTATGCGGCGACCGAGTTGCTCTTGCCCGGCGTCAATACGGGATAATACCGCGCCACATAGCAGAACTTTAAAAGTGCTCATCATTGGAAAACGTTCTTCGGGGCGAAAACTCTCAAGGATCTTACCGCTGTTGAGATCCAGTTCGATGTAACCCACTCGTGCACCCAACTGATCTTCAGCAT |

| | |
|---|---|
| | CTTTTACTTTCACCAGCGTTTCTGGGTGAGCAAAAACAGGAAGGCAAAATGCCGC<br>AAAAAAGGGAATAAGGGCGACACGGAAATGTTGAATACTCATACTCTTCCTTTTT<br>CAATATTATTGAAGCATTTATCAGGGTTATTGTCT |
| Plasmid pN (BspHI cut)[e] | CATGAGCGGATACATATTTGAATGTATTTAGAAAAATAAACAAATAGGGGTTCC<br>GCGCACATTTCCCCGAAAAGTGCCACCTGATGCGGTGTGAAATACCGCACAGAT<br>CGATCCCGCGAAATTAATACGACTCACTATAGGGGAATTGTGAGCGGATAACAA<br>TTCCCCTCTAGAAATAATTTTGTTTAACTTTAAGAAGGAGATATACCAAAGGCCA<br>GCAAAAGGCCAGGAACCGTAAAAAGGCCGCGTTGCTGGCGTTTTTCCATAGGCT<br>CCGCCCCCCTGACGAGCATCACAAAAATCGACGCTCAAGTCAGAGGTGGCGAAA<br>CCCGACAGGACTATAAAGATACCAGGCGTTTCCCCCTGGAAGCTCCCTCGTGCGC<br>TCTCCTGTTCCGACCCTGCCGCTTACCGGATACCTGTCCGCCTTTCTCCCTTCGGG<br>AAGCGTGGCGCTTTCTCATAGCTCACGCTGTAGGTATCTCAGTTCGGTGTAGGTC<br>GTTCGCTCCAAGCTGGGCTGTGTGCACGAACCCCCCGTTCAGCCCGACCGCTGC<br>GCCTTATCCGGTAACTATCGTCTTGAGTCCAACCCGGTAAGACACGACTTATCGC<br>CACTGGCAGCAGCCACTGGTAACAGGATTAGCAGAGCGAGGTATGTAGGCGGT<br>GCTACAGAGTTCTTGAAGTGGTGGCCTAACTACGGCTACACTAGAAGAACAGTA<br>TTTGGTATCTGCGCTCTGCTGAAGCCAGTTACCTTCGGAAAAAGAGTTGGTAGCT<br>CTTGATCCGGCAAACAAACCACCGCTGGTAGCGGTGGTTTTTTTGTTTGCAAGCA<br>GCAGATTACGCGCAGAAAAAAAGGATCTCAAGAAGATCCTTTGATCTTTTCTACG<br>GGGTCTGACGCTCAGTGGAACGAAAACTCACGTTAAGGGATTT**ATCGGGAGATC**<br>**TGGAGAAAAAAAAAAAAAAAGATACACATTGAGTGATTTC**GAGTAAACTTGGTC<br>TGACAGTTACCAATGCTTAATCAGTGAGGCACCTATCTCAGCGATCTGTCTATTTC<br>GTTCATCCATAGTTGCCTGACTCCCCGTCGTGTAGATAACTACGATACGGGAGGG<br>CTTACCATCTGGCCCCAGTGCTGCAATGATACCGCGAGACCCACGCTCACCGGCT<br>CCAGATTTATCAGCAATAAACCAGCCAGCCGGAAGGGCCGAGCGCAGAAGTGG<br>TCCTGCAACTTTATCCGCCTCCATCCAGTCTATTAATTGTTGCCGGGAAGCTAGAG<br>TAAGTAGTTCGCCAGTTAATAGTTTGCGCAACGTTGTTGCCATTGCTACAGGCAT<br>CGTGGTGTCACGCTCGTCGTTTGGTATGGCTTCATTCAGCTCCGGTTCCCAACGA<br>TCAAGGCGAGTTACATGATCCCCCATGTTGTGCAAAAAAGCGGTTAGCTCCTTCG<br>GTCCTCCGATCGTTGTCAGAAGTAAGTTGGCCGCAGTGTTATCACTCATGGTTAT<br>GGCAGCACTGCATAATTCTCTTACTGTCATGCCATCCGTAAGATGCTTTTCTGTGA<br>CTGGTGAGTACTCAACCAAGTCATTCTGAGAATAGTGTATGCGGCGACCGAGTT<br>GCTCTTGCCCGGCGTCAATACGGGATAATACCGCGCCACATAGCAGAACTTTAAA<br>AGTGCTCATCATTGGAAAACGTTCTTCGGGGCGAAAACTCTCAAGGATCTTACCG<br>CTGTTGAGATCCAGTTCGATGTAACCCACTCGTGCACCCAACTGATCTTCAGCAT<br>CTTTTACTTTCACCAGCGTTTCTGGGTGAGCAAAAACAGGAAGGCAAAATGCCGC<br>AAAAAAGGGAATAAGGGCGACACGGAAATGTTGAATACTCATACTCTTCCTTTTT<br>CAATATTATTGAAGCATTTATCAGGGTTATTGTCT |
| **C) Introns:** | |
| Mod. P.li.LSU.I2 intron RNA | GUGCGACAAGAAGUUCAGGAAGGGAUUGAGGUGAAAGUCCUCCUCCCGAAU<br>CGUUCAUGGGAGAGUCUAUCCAGACUUGCGUAGCGAGUAAUCGCUAGGUGA<br>GAAGCUCUGGAGACAAUGUACCUGCCCUUCAAUUGGAGGUGCCAGGGCUAG<br>GCUUUGUUCCUAUGGCUAGCGAAAGCGAAUACAGGCUAUUAGGCGUCGUGA<br>UCCUUAAACUCUUUCGACAUAAGGGAGAGAAAAGUAGAUGAGUUACCCUCC<br>CGCAACUGGGGUAACUGAACGAUCAAUAUCUACCGGCGUACACUGUGAGCC<br>UAAAGGAAACGAAUGAAAGUGUCCCUUCCUGGGAACUUGGUAAGCCCAAUA<br>GACUCCCUCUGGGAAACCAGAGGAGGAGUAAGAGCAAUCUUACUUAUCGUC<br>UAGUGGGUAAAAGACAUAAUAGGAAGCAAAUGCUUGGUUGUAACGAUCGA<br>GAUAGAAUCUGUUGAUUUAAGCUGAAAGGCUGCAGACUUAUUAAAUGGGU<br>GUUCUGCUUUCGGCAGCAGAAUGCGUGAGCCGUGUGCGAUGAAAGUCGCAA<br>GCACGGUUCUGAUGGGGGGAAAACGAGAGAUCGUCUACCUAUCCAACU |
| Mod. P.li.LSU.I2 $\psi_U$ intron RNA | GUGCGACAAGUAAGUUCAGGAAGGGAUUGAGGUGAAAGUCCUCCUCCCGAA<br>UCGUUCAUGGGAGAGUCUAUCCAGACUUGCGUAGCGAGUAAUCGCUAGGU<br>GAGAAGCUCUGGAGACAAUGUACCUGCCCUUCAAUUGGAGGUGCCAGGGCU<br>AGGCUUUGUUCCUAUGGCUAGCGAAAGCGAAUACAGGcUAUUAGGCGUCGU |

| | |
|---|---|
| | GAUCCUUAAACUCUUUCGACAUAAGGGAGAGaaaaGUAGAUGAGUUACCCUCCCGCAACUGGGGUAACUGAACGAUCAAUAUCUACCGGCGUACACUGUGAGCCUAAAGGAAACGAAUGAAAGUGUCCCUUCCUGGGAACUUGGUAAGCCCAAUAGACUCCCUCUGGGAAACCAGAGGAGGAGUAAGAGCAAUCUUACUUAUCGUCUAGUGGGUAAAAGACAUAAUAGGAAGCAAAUGCUUGGUUGUAACGAUCGAGAUAGAAUCUGUUGAUUUAAGCUGAAAGGCUGCAGACUUAUUAAAUGGGUGUUCUGCUUUCGGCAGCAGAAUGCGUGAGCCGUGUGCGAUGAAAGUCGCAAGCACGGUUCUGAUGGGGGGAAAACGAGAGAUCGUCUACCUAUCCAACU |
| C.i.SSU.I1 intron RNA | GUGCGACUUGUUAAGUUUUAACAAAAAUUGUAUAACGUUUAUUAAUGAUUAUACAUUGUAUUUCAUCUUACAAUAGCCUAAUUAGAUAUGCAUUUAGGGUAACUUUUUUGUAUAAAGCUCUAAUUAUAAGUGUAAAUACACUUUUAGGCUUCUUCUAUGGUUAGAGAAAUCGAACCAAUGUAAUUAAAACUUUGAUGUAUUAGGCAUUUAACGUGUCCUUGGUUAAAUGAAGAUGAACAUAAGUAUACAAAGUAAAAUUGGAACCUAAGGAAGAAUUGUUUUUGUUAAGAAACAAGGUAAUACCUAUAACUGGCUAAUAUAAAUUUGCAAGGUUUAUUGUAAAAUAAACUAUAGGUUAGAGGUAAAAGGAUAAUGUAAAAAGCGAAUGCAAUUCUGUAAUGGAAUUGAUAGGGUAUAUACCUAACUUGAAAGAGUGCUGACUUACAUAUAGAUGUUAUUUACGUuucgACGUAAAUAAUGUUUGAGCCGUAUGCUAUGAAAGUAGCACGUACGGUUCUAAGAGGGGGAAAGUCCGAGAGGACCUACCUAUCUCAAC |
| bI1 intron RNA | GUGAGACAAGUAUAAGUAUAUUAUUAUAAUAUCAUACCAUUAAAUAAAUUAUUUUAAUGAAAUGAUUAUGUUUAUAUAUAACAUAUACCUAAUUAGACAUGCAUUAUUAGUAAUAAUUUUGUAUGAAACUCUAAUAAUAAUAAUUAUUAUUAAUUAUUAAGGUAAGAUUCAUAUgGAUAGCGUAAGUCAAUCUAAUAUUAUAAAAUAUCGUAACAUAAACAAUAUUUUUUCUAUUAUUAAUUAAUAAAUAAUAAUAAAUAAAAAUAAUUAUAUGAGAAGUAAGAUAUUCAAUUCUGUCUAGAAUACAUAUAUAUACGUUAAUACUCAUCGGUAUAAAAUUAGAAUCCUAAGUGAAUUAUUGAAAGUAUAAUAAUAUAAACUUGGUAAGCCCAAUUAUUUCCAUAUAAUAUUAAUAUAAAUAUUAUAUGGUAGUUAUAUAUAAUAUUAUUAAAUAAAUAAUAAUAGAAAUUAUAAUAUAGAUAAGUGGGUAAAAGACUAUUGAAAAAGCUAAAGAUUAUAUGUAAUGUAUAAUAUAGAUCAAAUUAUUUAUAUAUUUUAAUAAAAAUAUAUUAAUAAUGGUUAAUAUUAUUAUUAAUUAAUUAAUUAAUUAAUUAAUAAUAAUAACGAAUAAAUGAUUAAUGUGAAAGCAUGCUAACUUCAAUAUAGGAUGAUUUAUAUAGUAUAUAAAUUGUUUGAGCUGUAUACUAUGAAAGUAGUACGUACAGUUCUGAGUGGGGGAAAAUUUGUAAAGAUCUACCUAUCACAAU |
| **D) PCR and RT-PCR primers:** | |
| P.li.LSU.I2 template FWD | GCGTAATACGACTCACTATAGGAGCGA |
| P.li.LSU.I2 & C.i.SSU.I1 template REV | AATCTTAGGACCCTCAGTGAACGC |
| C.i.SSU.I1 template FWD | GCGTAATACGACTCACTATAGGAGCGAA |
| bI1 template FWD | GCGTAATACGACTCACTATAG |
| bI1 template REV | AGCGGATAACAATTTCACACAGG |
| RS plasmid RT primer | GAAGGGCAGGTACATTGTCT |
| RS plasmid RT-PCR FWD | TGGCCTAACTACGGCTACACTA |
| RS plasmid RT-PCR REV | CAGAGCTTCTCACCTAGCGATTAC |
| ssDNA 3'SS RT primer[f] | GAAATCACTCAATGTGTAC |
| ssDNA 3'SS RT-PCR FWD | GGCTGCAGACTTATTAAATG |
| ssDNA 3'SS RT-PCR REV | GAAATCACTCAATGTGTAC |
| ssDNA 5'SS RT primer | GATCACGACGCCTAATAG |
| ssDNA 5'SS RT-PCR FWD | GCGTAATACGACTCACTATAG |
| ssDNA 5'SS RT-PCR REV | GATCACGACGCCTAATAG |

[a] The T7 promoter is in italics. The sequence corresponding to the intron is in boldface. Underlined portions of the sequence correspond to the deviations from the crystallization construct from Robart *et al.* 2014 (9).

[b] The red “T” denotes the inserted base.

[c] Fluorescent labels are denoted in the IDT format. /5Cy5/ is 5’-Cy5. /36-FAM/ is 3’-6FAM. /iCy5/ is internal Cy5. /iFluorT/ is internal Fluorescein dT.

[d] The underlined bases denote parts of the mod. P.li.LSU.I2 IBSs. Blue is IBS2a, red is IBS2, green is IBS1, orange is IBS3.

[e] Boldface denotes either the insert carrying the intron IBSs, or the control insert lacking the IBSs.

[f] Only the underlined region is complementary to the short 3’exon of the DNA RS sub. 2.

### Supplementary references

1. Monachello,D., Lauraine,M., Gillot,S., Michel,F. and Costa,M. (2021) A new RNA–DNA interaction required for integration of group II intron retrotransposons into DNA targets. *Nucleic Acids Res.*, **49**, 12394–12410.

2. Nagy,V., Pirakitikulr,N., Zhou,K.I., Chillón,I., Luo,J. and Pyle,A.M. (2013) Predicted group II intron lineages E and F comprise catalytically active ribozymes. *RNA*, **19**, 1266–78.

3. Mörl,M., Niemer,I. and Schmelzer,C. (1992) New reactions catalyzed by a group II intron ribozyme with RNA and DNA substrates. *Cell*, **70**, 803–810.

4. Toor,N. and Zimmerly,S. (2002) Identification of a family of group II introns encoding LAGLIDADG ORFs typical of group I introns. *RNA*, **8**, 1373–1377.

5. Mullineux,S.-T., Costa,M., Bassi,G.S., Michel,F. and Hausner,G. (2010) A group II intron encodes a functional LAGLIDADG homing endonuclease and self-splices under moderate temperature and ionic conditions. *RNA*, **16**, 1818–1831.

6. Liu,T. and Pyle,A.M. (2021) Discovery of highly reactive self-splicing group II introns within the mitochondrial genomes of human pathogenic fungi. *Nucleic Acids Res.*, **49**, 12422–12432.

7. Skilandat,M. and Sigel,R.K.O. (2014) The Role of Mg(II) in DNA Cleavage Site Recognition in Group II Intron Ribozymes. *J. Biol. Chem.*, **289**, 20650–20663.

8. Steffen,F.D., Khier,M., Kowerko,D., Cunha,R.A., Börner,R. and Sigel,R.K.O. (2020) Metal ions and sugar puckering balance single-molecule kinetic heterogeneity in RNA and DNA tertiary contacts. *Nat. Commun.*, **11**, 104.

9. Robart,A.R., Chan,R.T., Peters,J.K., Rajashankar,K.R. and Toor,N. (2014) Crystal structure of a eukaryotic group II intron lariat. *Nature*, **514**, 193–197.
